# An uncanonical transcription factor-DREB2B regulates seed vigor negatively through ABA pathway

**DOI:** 10.1101/2020.12.09.418798

**Authors:** Faiza Ali, Zhenzhen Wei, Yonghui Li, Lei Gan, Zuoren Yang, Fuguang Li, Zhi Wang

**Affiliations:** Zhengzhou Research Base, State Key Laboratory of Cotton Biology, Zhengzhou University, Zhengzhou, 450001, China; State Key Laboratory of Cotton Biology, Key Laboratory of Biological and Genetic Breeding of Cotton, Institute of Cotton Research, Chinese Academy of Agricultural Sciences, Anyang, 455000, China

**Keywords:** DREB transcription factor, seed germination, seed vigor, tetrazolium, Abscisic acid, Controlled Deterioration Test

## Abstract

Seed vigor is an important trait for ecology, agronomy, and economy and varies with different plant species and environmental conditions. Dehydration-Responsive Element-Binding Protein 2B (DREB2B), a subgroup of the DREB transcription factor family, is well-known in drought resistance. However, the role of *DREB2B* in the regulation of seed vigor has not been identified. Here, we found that *DREB2B* is a negative regulator of seed vigor by ABA-mediated pathway in Arabidopsis with loss of function mutant and over-expressed transgenic lines. Furthermore, *DREB2B* showed epistatic and parallel to *ABI3* simultaneously in seed vigor regulation by genetic and molecular approaches. *DREB2B* homolog gene (*GhDREB2B-A09*) was also identified in cotton. The expression analysis indicated that transcripts of *DREB2B* were higher in mature dry seed, and the transgenic plants showed the conservative roles of *DREB2B* in Arabidopsis and cotton. In addition, we identified that DREB2B interacted with RADICAL-INDUCED CELL DEATH1 (RCD1) to involve seed vigor regulation together in *Arabidopsis* and cotton with BiFC experiment and mutant phenotypic analysis. Collectively it is concluded that DREB2B interacting with RCD1 or SRO1 function at upstream of and synergistic with *ABI3* to regulate seed vigor negatively in *Arabidopsis* and cotton, which provides novel knowledge in the seed development study.

**Highlights:** DREB2B transcription is seed specific and a negative regulator of seed vigor by ABA-mediated pathway, which interacts with RCD1s, and functions synergistically with ABI3 to affecet seed germination and vigor in Arabidopsis and cotton.

## Introduction

Ecologically, the seed is an essential tissue for their survival, diffusion, and initiation of the succeeding generation in plant species (Koornneef *et al*., 2002). Seed longevity determined by seed vigor is a quantitative trait defined as the period of seed dry storage in which the seed does not lose its viability and remains capable to germinate after maturation (Angelovici *et al*., 2010; Bartee and Krieg, 1974). It is gradually established during seed development and is necessary for plant survival during unfavorable conditions. However, it decreases with the seed storage life span in dry storage conditions (Abdelmagid and Osman, 1975; Jing-bao *et al*., 2011) and might be affected by temperature and moisture content in many species (Bailly *et al*., 2004). Therefore besides natural aging, artificial accelerated aging in which seeds are stored at a higher temperature and moisture conditions, is used to determine seed longevity (Oracz *et al*., 2007; Yamaguchi *et al*., 2007) and known as “Controlled Deterioration Treatment (CDT)” or “Accelerated Aging” in the field of seed longevity search (Chen *et al*., 2016; Hay *et al*., 2019).

Abscisic acid (ABA) shows significant functions in plant development including seed dormancy and germination, primary root growth, seedling development, reproductive growth, and response to abiotic stresses (Lovegrove and Hooley, 2000; Razem *et al*., 2006; Shu *et al*., 2013). In a study, ABA also showed an important role in seed vigor and longevity (Dekkers *et al*., 2016). In addition, mutations in many *Arabidopsis* genes involved in ABA biosynthesis and signaling pathway controlling seed maturation and dormancy led to a reduction in seed vigor and viability such as *lec1-3, aba1-5, abi3-1, abi3-5*, and *abi3-7* mutants, which showed significantly shorter longevity and lower vigor under ambient storage conditions (Cantoro *et al*., 2013; Hu *et al*., 2017; Liu *et al*., 2013; Ravindran *et al*., 2017; Sugliani *et al*., 2009). Moreover, *ABI3* plays a central role upstream of heat shock transcription factor HSFA9 to involve seed longevity and vigor control in *Arabidopsis* (Kotak *et al*., 2007; Sano *et al*., 2016; Tejedor-Cano *et al*., 2010).

Dehydration-Responsive Element-Binding Protein 2B (DREB2B) is a member of the DREB2 transcription factor family belonging to the subfamily of Ethylene-responsive element-binding proteins (EREBP) TF family (Okamuro *et al*., 1997; Sakuma *et al*., 2002; Weigel, 1995). *DREB2B* encodes *DRE/CRT* (one of the major *cis*-acting elements function in ABA-responsive or non-responsive gene expression during abiotic stresses) -binding protein *DRE/CRT* (Liu *et al*., 1998; Nakashima and Yamaguchi-Shinozaki, 2010). The *DREB2B* gene was identified as a key transcription factor that functions particularly in dehydration and heat stress response (Liu *et al*., 1998). However, the role of DREB2B related to seed vigor and longevity is unknown. Here, the higher seed germination rate, stronger seed vigor, and longevity, and reduced germination sensitivity to ABA of *dreb2b* knockout mutant illustrated that *DREB2B* is a negative regulator in seed vigor and longevity by ABA-mediated pathway. Genetic analysis showed that DREB2B function epistatic and synergistic with ABI3 to involve seed vigor through different pathways. Besides, it is suggested that DREB2B interacted with RCD1 family proteins to form a complex to negatively control seed vigor in *Arabidopsis* and cotton.

## Materials and Methods

### Genetic materials and growth conditions

The *dreb2b* (SALK_102687C), *abi3-16* (SALK_023411C), *rcd1-3* (SALK_116432), *sro1-1* (SALK_074525), *sro1-2* (SALK_126383) mutants were got from the Arabidopsis Biological Resource Center (ABRC) and belong to Columbia-0 (Col-0) background as well as Col-0 wild type (WT) was used as a control for all experiments (Supplementary Table S1). *rcd1-3*, and *sro1-1* mutants were identified as before (Jaspers *et al*., 2009). *dreb2b/abi3-16* double mutant was acquired by crossing the single homozygous *dreb2b* and *abi3-16* mutants. All homozygous single and double mutant lines were isolated by PCR-based screening using gene-specific (LP/RP) and T-DNA (BP/RP) primers of respective mutant lines obtained from Salk Institute Genomic Analysis Laboratory (SIGnAL) database (Supplementary Table S2). PCR was done with 35 cycles stand on the primers’ annealing temperature.

WT, mutants, and double mutant seeds were surface sterilized for 15-mints in 10% bleach and washed at least five times with double distilled water. Sterilized seeds were kept for 72 h at 4°C in dark for stratification and sown on ½MS medium at 22°C, with 16h light/8h dark photoperiod. 15-days old seedlings were moved to soil and grown in a growth chamber at 22°C. Seeds used in experiments were reaped and placed under dry conditions at 25°C.

### Germination analysis

To determine the germination ratio, seed vigor, and sensitivity to ABA and PAC, seeds were collected from WT and mutant plants of different genotypes grown simultaneously and stored under the same conditions. For germination and seed vigor tests, seeds from all genotypes were spread on filter paper moistened with distilled water in 5cm Petri dishes; while, for ABA and PAC sensitivity, all lines were soaked on filter paper with mock solutions or solutions enriched with ABA and PAC concentration in Petri dishes, and all germination analysis was performed as described previously (Bentsink *et al*., 2006).

### Controlled Deterioration Test (CDT)

CDT was performed as described previously with minor modifications (Liu *et al*., 2015; Yamaguchi *et al*., 2007). Mutants and WT seeds were harvested at the same time, dried, and then stored under the same conditions two weeks earlier to the experiment. The seeds were stored in a closed glass container saturated with KCl solution to provide 82% relative humidity (RH) for 3-days at room temperature for equilibration. After equilibration, the seeds were stored at 42°C with 82% relative humidity (RH) in a temperature and humidity controlled incubator for 3-, 5- and 7-days. After CDT treatment seeds were dried at room temperature for two days and tested for germination as described previously or other determination (Bentsink *et al*., 2006).

### Tetrazolium assay for seed viability

Tetrazolium assay was performed as described in an online published protocol with little modification (Rajjou *et al*., 2008; Verma *et al*., 2013). With or without CDT treated seeds were initially surface sterilized with 10% hypochlorous acid (having 0.1% Triton X-100) for 15 minutes and washed five times with sterilized distilled water. After that seeds were socked in 1% tetrazolium solution (pH-7.0) at 30°C in darkness for 2 days. After staining seeds were washed three times with sterilized distilled water. Seeds viability was determined by investigating the color intensity and staining pattern. Viable seeds turned to red, while non-viable or dead seeds stayed unstained.

### Gene identification and bioinformatics analysis

For the identification of *DREB2B* gene homolog in cotton (*G. hirsutum*), the *Arabidopsis DREB2B* (AT3G11020.1) gene was used as query and its homolog gene was identified in *G. hirsutum* (NAU, version 1.1) via BLAST search, and protein sequences were further verified by hidden Markov model (HMM) as described previously (Faiza *et al*., 2019; QANMBER *et al*., 2018). As a result, *Gh_A09G2423* was identified as its closest homolog which was named *GhDREB2B-A09*. For alignment, firstly, multiple sequence alignment was created in Clustal X 2.0 (http://www.clustal.org/clustal2/) and alignment was visualized by online tool ESPript (http://espript.ibcp.fr/ESPript/cgi-bin/ESPript.cgi). The phylogenetic tree was created through MEGA 7.0 (Kumar *et al*., 2016) using the neighbor-joining (NJ) method with 1000 bootstrap replicates with 50% cutoff values as described previously (Qanmber *et al*., 2019c). Domain prediction was carried out by Interproscan 63.0 (http://www.ebi.ac.uk/InterProScan/) and SMART (http://smart.embl-heidelberg.de/) search. For promoter *cis*-element analysis, 2kb upstream of the start codon was subjected to PlantCARE (Ghaderi-Far *et al*., 2012) database and the figure was generated using TBtools (Forman and Jensen, 1965).

### Expression analysis

To determine gene expression in different developmental stages, tissues such as root, stem, leaf, flower, siliques at 3, 6, 9, 12, 15, 18 days after pollination (DAP), and mature seeds, were collected from *Arabidopsis* plant grown under normal growth conditions. To collect siliques at different developmental stages, blooming flowers were first marked by tying with cotton threads on the day of pollination, and then siliques were collected on a required day after pollination. Further, cotton tissues were also collected at different developmental stages including root, stem, leaf, flower, ovules of −2, 0, 3, 5, 10, 15, 20, and 25 DPA (day post-anthesis) as well as fiber tissues of 1, 10, 15, and 20 DPA, and mature seed from ZM24 (CCR124) cotton plant, grown under field conditions in Zhengzhou China (Yang *et al*., 2019; Yang *et al*., 2014). To collect imbibed seeds, seeds were imbibed in double-distilled water and kept at room temperature, and collected after different hour intervals (2, 6, 12, 18, and 24 h).

The total RNA from all collected tissues was isolated using RNAprep Pure Plant Kit (TIANGEN, Beijing, China), as per the manufacturer’s instructions. For complementary strand (cDNA) synthesis and PCR amplification, Prime Script® RT reagent kit (Takara, Dalian, China) and SYBR Premix Ex Taq™ II (Takara) was used (Qanmber *et al*., 2019a). *ACTIN2 (At3G18780*.*1)* (Wang *et al*., 2013) and *GhHis3* (AF024716) (Wan *et al*., 2016) were used as a control for *Arabidopsis* and cotton respectively and PCR was performed with three independent biological replicates. The relative expression values were calculated and used to draw the figures as described before (Ali *et al*., 2020; Qanmber *et al*., 2019b).

To evaluate the mRNA level of ABA biosynthesis, signaling and catabolism genes in *dreb2b* mutant dry seeds and mRNA level of *DREB2B, ABI3, RCD1*, and *SRO1* in *dreb2b, abi3-16, rcd1-3, sro1-1, dreb2b/abi3-16, snl1snl2-1* mutants, the RNA was extracted from freshly harvested WT and mutant seeds as described above. The primers used for qRT-PCR in this study were enlisted in Supplementary Table S2.

### Plasmid construction

For plasmids construction, the full-length coding region and 1.6 and 1.8 promoter fragments of *DREB2B* and *GhDREB2B-A09* were amplified from *Arabidopsis* and ZM24 seeds cDNA and DNA respectively using high-fidelity DNA polymerase. The amplified CDS were then fused with pCAMBIA-2300 containing GFP-tag for over-expression and complementary lines as well as subcellular localization analysis, under the control of 35S constitutive promoter; By gateway technology, the CDS were also transferred to pAS2-attR containing GAL4 binding domain for transcriptional activity. The promoters were fused into pBI121 (K, 2007) containing the GUS gene for promoter-GUS activity analysis. The primers used for amplification of genes coding regions and promoter were enlisted in supplementary Table S2.

### Transformation

The fusion genes for over-expression, promoter-GUS activity, and complementation were then individually introduced into Col-0 and *dreb2b* mutant plants respectively via the floral dip method (Zheng *et al*., 2020) with *Agrobacterium tumefaciens* strain (GV3101). After that harvested seeds were surface sterilized as described above. Positive lines were selected on ½MS medium plates holding kanamycin (50 mg/L) incubated under 16 h / light and 8 h / dark cycle at 22°C. The 3:1 segregating transformants lines were selected on MS medium and further verified by PCR. T3 or T4 homozygous transgenic plants were used for analyses.

### Transcriptional activity assay

To examine the transcriptional activity, fusion constructs of *pBD-DREB2B* and *pBD-GhDREB2B-A09* were individually introduced into Y2H-Gold yeast cells and spread on a medium plate lacking Trp, incubated at 28°C for 2-4 days. The positive colonies were cultured in an SD/-Trp liquid medium to OD600 of approximately 1, then the culture was diluted 10 to 100 times with fresh medium and grown further on three selected medium plates such as SD/-Trp, SD/-Trp-His, and SD/-Trp-His+x-α-gal at 28°C for 2-4 days. The transcriptional activity on three selected medium plates was observed and images were captured with a digital camera (Canon).

For β-galactosidase assay constructs were processed according to the instructions of the β-galactosidase assay kit provided by Clontech. For each construct, five colonies were assayed and β-galactosidase activity was expressed in miller units.

### Subcellular localization and GUS staining

For protein localization, tobacco *(Nicotiana benthamiana)* leaves were co-infiltrated with the *Agrobacterium* strains (GV3101) containing the *35S:DREB2B-GFP* and *35S:GhDREB2B-A09-GFP* construct along with p19 (Smith, 1995). The infiltrated plants were kept in dark for 24 h and then in light for 48 h. The protein localization images from tobacco epidermal cells were obtained with a fluorescence microscope (Olympus) after 72 h of infiltration.

To verify the transient expression, protein localization was determined in root and radical cells of homozygous *35S:DREB2B* and *35S:GhDREB2B-A09* lines at T4 generation. 15-days old seedlings and radical cells after 24 h imbibitions were used to detect the GFP fluorescence with 1µg/mL–14’, 6-diamidino-2-phenylindole (DAPI) (cell nuclei staining dye) (Sigma-Aldrich), and images from the root and radical cells were obtained with a fluorescence microscope (Olympus).

To determine the tissue-specific localization, freshly collected seedlings and tissues from T4 transgenic plants of pBI121-promoter were incubated into GUS solution (Yuan Ye, Shang Hai) at 37°C overnight according to the manufacturer’s instructions. After staining, samples were rinsed 3 to 4 times with 70% ethanol and images were captured with a stereo-microscope.

### Bimolecular fluorescence complementation (BiFC)

Firstly, the potential interaction factors of DREB2B and GhDREB2B-A09 were predicted by the online prediction tools Arabidopsis eFP Browser (http://bar.utoronto.ca/efp/cgi-bin/efpWeb.cgi) and ccNET (http://structuralbiology.cau.edu.cn/gossypium/cytoscape/network.php) (Turley and Chapman, 2009) respectively. The predicted interactions between DREB2B and RCD1 or SRO1 were further verified by BiFC. First, the full-length coding regions of *DREB2B, RCD1, SRO1, DREB2BL-A09, GhRCD1L-A05, GhRCD1L-D05, GhRCD1L-A08, GhRCD1L-D12*, and *GhRCD1L-A12* were amplified from *Arabidopsis* and cotton cDNA (Supplementary Table S2). The amplified fragments were then cloned into vectors *pXY104* and *pXY106*, prepared *nYFP-DREB2B, nYFP-DREB2BL-A09, cYFP-RCD1, cYFP-SRO1, cYFP-GhRCD1L-A05, cYFP-GhRCD1L-D05, cYFP-GhRCD1L-A08, cYFP-GhRCD1L-D12*, and *cYFP-GhRCD1L-A12* constructs were transformed into GV3101 and were co-infiltrated into 5-6 weeks old tobacco leaves as described above. The YFP fluorescence signals were perceived with a fluorescence microscope (Olympus).

## Results

### DREB2B negatively regulates seed longevity and vigor

Previously, the expression of the *DREB2B* gene was down-regulated in an *snl1snl2-1* double mutant which showed clearly decreased seed dormancy and longevity phenotypes (Wang *et al*., 2013), suggesting that *DREB2B* might have functions in seed related traits. To investigate the role of *DREB2B* in seed related traits, loss of function mutant *dreb2b* with a T-DNA insertion at 513bp in exon was identified as a null mutant (Supplementary Fig. S1A). Firstly, we analyzed the germination ratio of freshly harvested seeds in both WT and *dreb2b* plants after different week’s intervals. *dreb2b* showed a slightly higher seed germination rate to that of WT within a short storage period (room temperature) (Supplementary Fig. S1B), indicating the potential role of *DREB2B* in seed vigor. Further, the percentage of seed germination was measured after 6, 12, 18, and 24 months interval during a longer period of dry storage and showed that *dreb2b* showed significantly higher (63%) seed germination than that of WT (26%) after 24 months of dry storage (Fig. 1A), which indicated that *DREB2B* might negatively regulate seed longevity. Additionally, the germination speed of freshly harvested seeds was also examined after two weeks of dry storage. After 96 h imbibition following stratification, *dreb2b* showed 60% germination while WT showed 16%. Although, 95% germination of *dreb2b* and WT was observed at 168 h imbibition respectively (Fig. 1B, C), signifying the quicker seed germination in *dreb2b* and negative role of *DREB2B* in seed vigor. Moreover, the role of *DREB2B* was further confirmed by *DREB2B* complementary assays in *35S:DREB2B/dreb2b* lines. The seeds from two independent complementary lines exhibited a similar germination speed to that of WT after stratification (Supplementary Fig. S1C, D), which verified the role of *DREB2B* in seed vigor.

**Fig. 1.**
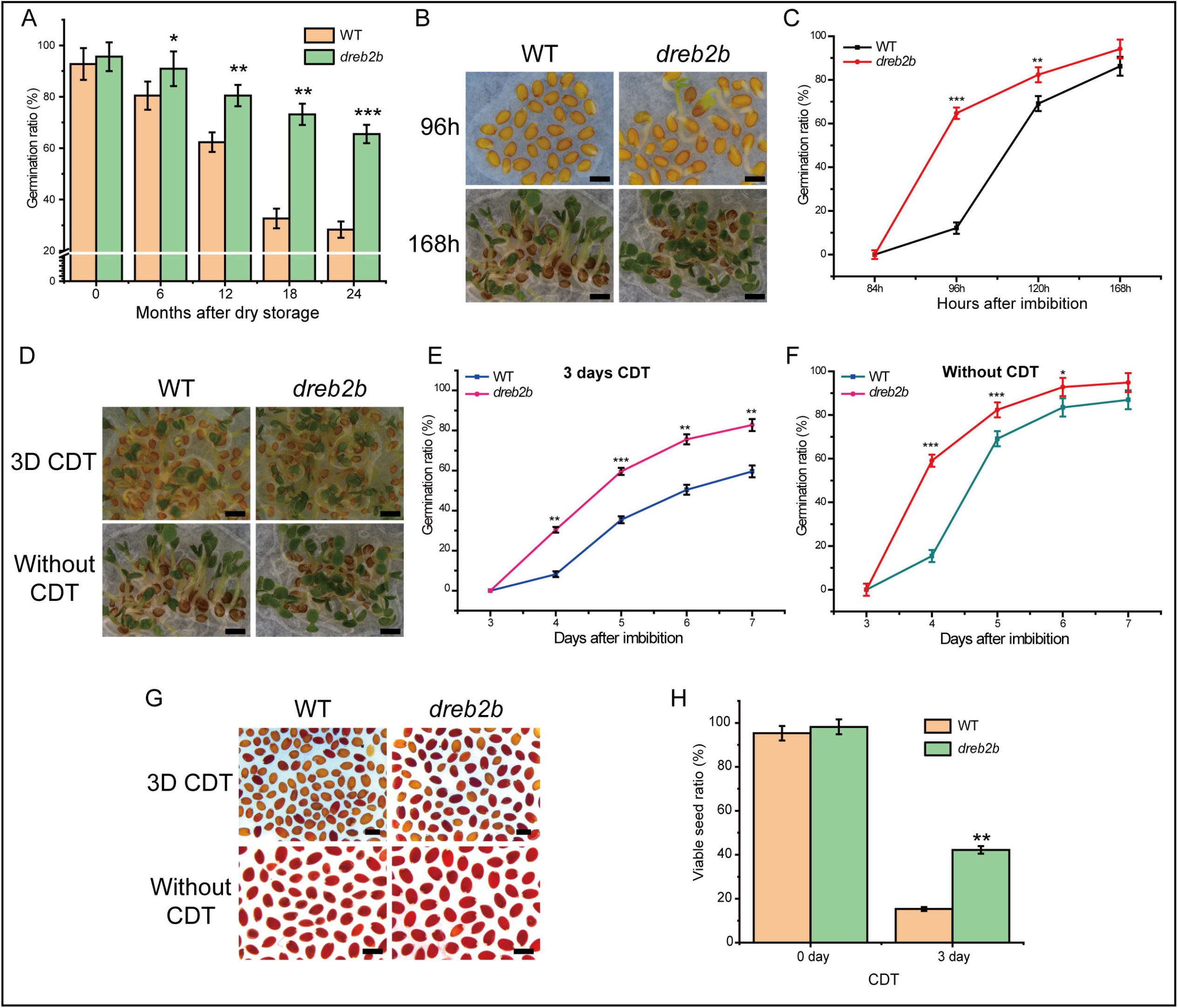
Phenotypic analysis of *dreb2b* in seed germination and vigor (A) Seed germination and vigor analysis of *dreb2b* mutant and WT after different months of dry storage at room temperature. The seed germination rate of *dreb2b* mutant and WT was calculated at 7 days after imbibition followed stratification. (B) Germinating seeds images of WT and *dreb2b* were taken at 84 and 168 hours after imbibition followed stratification. (C) *dreb2b* mutant and WT seed germination speed after different hour’s interval was calculated from 2 weeks old seeds harvested at the same time. (D) Seed germination images of WT and *dreb2b* with 3 day CDT or without CDT were taken at 7 days after imbibition with stereo-microscope. (E, F) WT and *dreb2b* seed germination rate treated with 3 days CDT or without CDT treatment was calculated with different day’s interval after imbibition. (G) Tetrazolium staining photos of WT and dreb2b mutant seeds. Tetrazolium staining was performed with and without CDT treated seeds. (H) Viable seed ratios of WT and *dreb2b* were evaluated by examining the tetrazolium staining pattern and color intensity. (B, D, G) For each genotype, 80-100 seeds were used. Bars, 500µm. (C, E, F) Experiments were performed in three replicates and germination ratio was measured from averages of three replicates with 100 seeds per replicate from independent lines of each genotype. Error bars indicate SD based on three biological replicates. Asterisks over the bars indicate a significance level among means determined by the student’s T-test (*P□ 0.05, **P□ 0.01, ***P□ 0.001).

Furthermore, the *dreb2b* seed longevity was analyzed by CDT too. The germination ratios were 59% and 83% for WT and *dreb2b* respectively after 3-days CDT treatments. In contrast, the germination ratio of untreated seeds were 85% and 95% for WT and *dreb2b* respectively (Fig. 1D-F). In addition, tetrazolium staining displayed that seeds of WT and *dreb2b* showed 100% viability without CDT treatment, whereas, after 3-days CDT treatment, a higher number of alive seeds in *dreb2b* (42%) mutant than WT (15%) was observed (Fig. 1G, H). CDT treatment and tetrazolium staining further confirmed the negative role of *DREB2B* in seed vigor and longevity.

### DREB2B regulates seed vigor and germination dependent on ABA pathway

Abscisic acid (ABA) and gibberellins (GA) are pivotal phytohormones in seed germination and vigor regulation. To investigate whether *DREB2B* functions in ABA or GA dependent pathway to direct seed vigor and germination, *dreb2b* seed germination was determined by applying exogenous ABA and PAC (Pactobutrazol, an inhibitor for GA synthesis). As a result, *dreb2b* showed increased germination compared to WT with different ABA concentrations (Fig. 2A) suggesting the hyposensitivity of *dreb2b* for ABA. However, after PAC treatment *dreb2b* showed similar germination to that of control (WT) (Supplementary Fig. S2A). These results pointed out that *DREB2B* may be involved in the ABA-mediated pathway to negatively regulate seed germination and vigor.

**Fig. 2:**
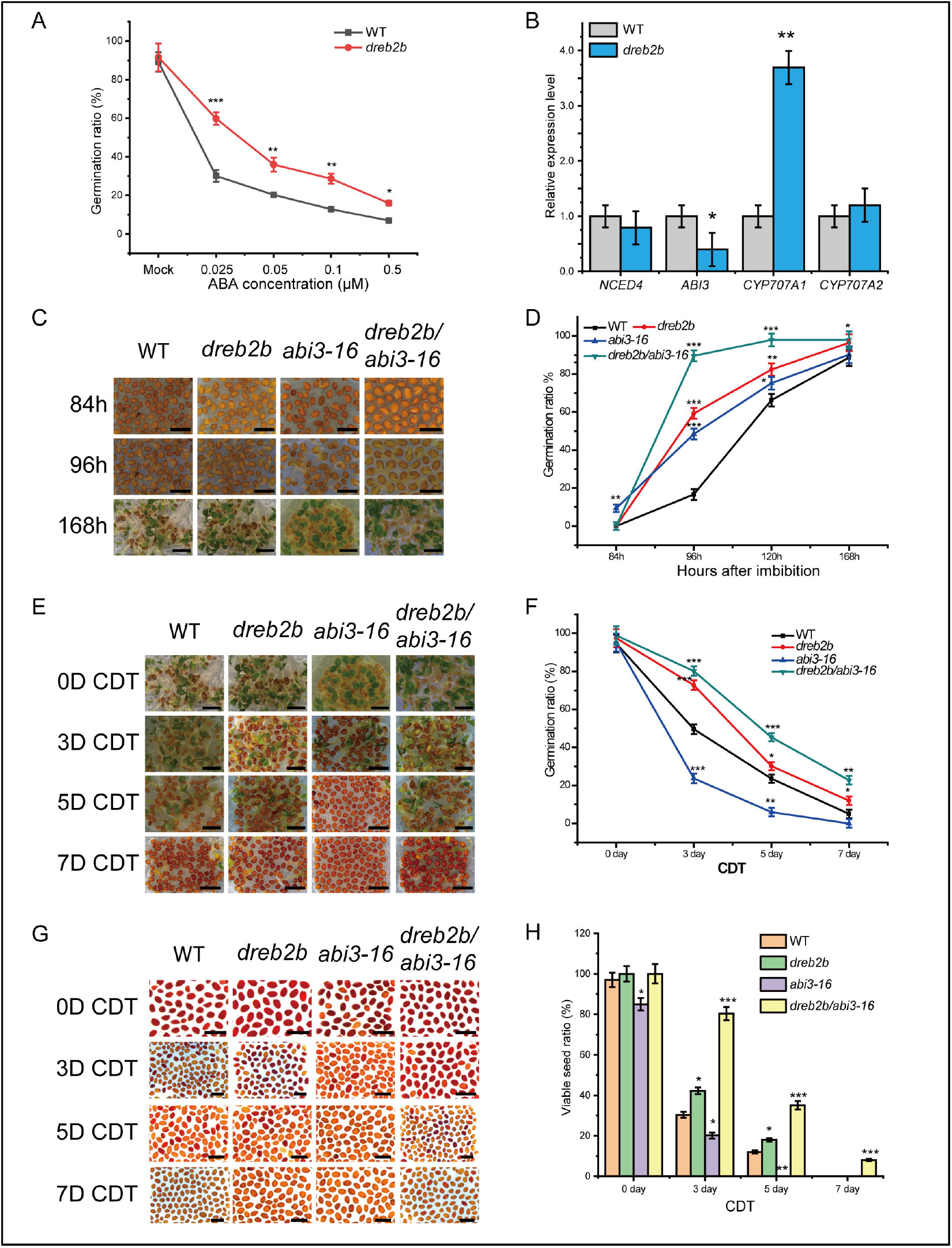
*DREB2B* regulated the seed germination and vigor through the ABA pathway. (A) Seed germination test of WT and *dreb2b* under ABA treatments. Freshly harvested seeds were soaked in distilled water with (0.025, 0.05, 0.1, and 0.5) or without ABA (Mock). The germination rate was counted at 7 days after imbibition followed stratification. (B) mRNA level of genes involved in ABA pathway in freshly harvested *dreb2b* mutant and WT dry seeds. Error bars indicate SD based on three biological replicates. Asterisks over the bars indicate a significance level among means determined by the student’s T-test (*P□ 0.05, **P□ 0.01, ***P□ 0.001). (C) Microscopic images of WT, *dreb2b, abi3-16* mutants, and *dreb2b/abi3-16* double mutant seeds showing germination speed were taken at 84, 96 and 168 hours after imbibition followed stratification. (D) WT, *dreb2b, abi3-16* mutants, and *dreb2b/abi3-16* double mutant seed germination after different hour intervals were observed and calculated from fresh seeds harvested 2 weeks before the experiment. (E) Images of WT, *dreb2b, abi3-16* mutants, and *dreb2b/abi3-16* double mutant with 0, 3, 5, and 7 days CDT were taken at 7 days after imbibition with stereo-microscope. (F) WT, *dreb2b, abi3-16*, and *dreb2b/abi3-16* double mutant seeds were treated with 0, 3, 5, and 7 days CDT, and the germination ratio of treated seeds was calculated at 0, 3, 5, and 7 days after imbibition. (G) Photos of WT, dreb2b, abi3-16 mutants, and dreb2b/abi3-16 double mutant seeds stained by tetrazolium after 0, 3, 5, and 7 days CDT treatments were taken with stereo-microscope. For each genotype, 80-100 seeds were used. Bars, 500µm. (H) Viable seed ratio of WT, dreb2b, abi3-16 mutants, and dreb2b/abi3-16 double mutant treated with 0, 3, 5, and 7 days CDT were calculated by examining the tetrazolium staining pattern and color intensity. (C, E) For each genotype, 80-100 seeds were used. Bars, 1mm. (A, D, F, H) Experiments were performed in three replicates and germination ratio was measured from averages of three replicates with 100 seeds per replicate from independent lines of each genotype. Error bars indicate SD based on three biological replicates. Asterisks over the bars indicate a significance level among means determined by the student’s T-test (*P□ 0.05, **P□ 0.01, ***P□ 0.001).

Many genetic studies indicated that the ABA metabolism and signaling genes are important in the regulation of seed germination (Finkelstein *et al*., 2002; Matilla *et al*., 2015; Nambara *et al*., 2010; Sun, 2008). Thus, the expressions of key genes such as *NCED4, ABI3, CYP707A1*, and *CYP707A2* were analyzed in the *dreb2b* mutant. The result showed that the expressions of *ABI3* and *CYP707A1* were significantly down-regulated and up-regulated in *dreb2b* mutant respectively (Fig. 2B) indicating that *DREB2B* might regulate seed germination and vigor by regulating the *ABI3* involved ABA signaling positively and *CYP707A1* mediated ABA hydrolysis negatively.

### DREB2B functions upstream of and synergistically with ABI3 to involve seed vigor regulation

*ABI3* is a key regulator in the ABA signaling pathway, positively controlling the seed longevity and the seed dormancy (Finkelstein *et al*., 2008; Hu *et al*., 2017; Yamaguchi *et al*., 2007). Here, we identified the clearly decreased transcription of *ABI3* in *dreb2b* (Fig. 2B). Moreover, the *DREB2B* transcript level in *abi3-16* mutant seeds was almost the same as in WT (Supplementary Fig S2B). To further explore the relationship between *DREB2B* and *ABI3, dreb2b/abi3-16* double mutant was generated and the expression of both genes (*DREB2B* and *ABI3)* was assessed by qRT-PCR. Significantly down-regulated expression of *DREB2B* and *ABI3* in *dreb2b/abi3-16* elucidating that *dreb2b/abi3-16* is a loss of function double mutant of both genes (Supplementary Fig S2C). Additionally, the up-regulated expression of *DREB2B* with different ABA concentrations indicated that *DREB2B* is induced more significantly than ABI3 by ABA (Supplementary Fig S2D). To deeply investigate the correlation between *DREB2B* and *ABI3* in the regulation of seed germination and vigor, germination speed was analyzed in freshly harvested seeds for *dreb2b/abi3-16* double mutant, *dreb2b, abi3-16* single mutants, and WT. Similar early germination was observed in both single mutants *(dreb2b* and *abi3-16)*, while *dreb2b/abi3-16* showed more quick germination (89%) than single mutants and WT (Fig. 2C, D), which indicated that DREB2B may be negatively involved in seed germination synergistically with ABI3. *dreb2b/abi3-16* seed longevity and viability was further determined by different days (0, 3, 5, and 7) CDT treatments. After 3, 5, and 7-days CDT, germination and viability were significantly reduced in all genotypes, and the *abi3-16* was arrested most severely, whose germination decreased to 23% just after 3-days CDT and completely loosed seed viability after 5-days CDT supporting the previous report (Clerkx *et al*., 2004; Sugliani *et al*., 2009); *dreb2b* was less affected by CDT compared to *abi3-16* and WT and loosed its viability until 7-days CDT same as WT. Whereas, germination of *dreb2b/abi3-16* were 80%, 45%, and 22% after 3, 5, and 7-days CDT treatments, respectively; in addition, it showed 12% viable seeds even after 7-days CDT which is significantly higher than other lines (Fig. 2E-H).

Summarily, the *abi3-16* mutation in *dreb2b* increased the seed germination and vigor significantly, suggest that ABI3 and DREB2B are involved in seed germination and vigor additively and synergistically. Combing the seed germination and gene expression analysis, it indicated that DREB2B might function upstream of and synergistic with ABI3 in seed germination and vigor regulation through different pathways.

### *DREB2B* transcription is seed-specific in Arabidopsis and cotton

Cotton is a kind of important cash crop, the uniform and quick germination of cotton seed are vital for its quality and yield. So, *DREB2B* homolog genes were identified in *G. arboreum, G. hirsutum, G. barbadense, G. raimondii, Z. may*, and *O. sativa* to explore the potential functional genes in seed vigor regulation of cotton. A total of ten *DREB2B* homologous genes (Six in cotton, two in maize, and two in rice) were identified and a phylogenetic tree was constructed with a maximum similarity matrix. The result illustrated the maximum similarity of *DREB2B* with *Gh_A09G2423*, which was studied in our research and renamed as *GhDREB2B-A09* (Fig. 3A). Further protein sequence analysis indicated that both *Arabidopsis* and cotton DREB2Bs (DREB2B and GhDREB2B-A09 respectively) show the low identity of amino acids with conserved AP2 domains and low complexity regions (LC) (Supplementary Fig. S3A, B).

**Fig. 3:**
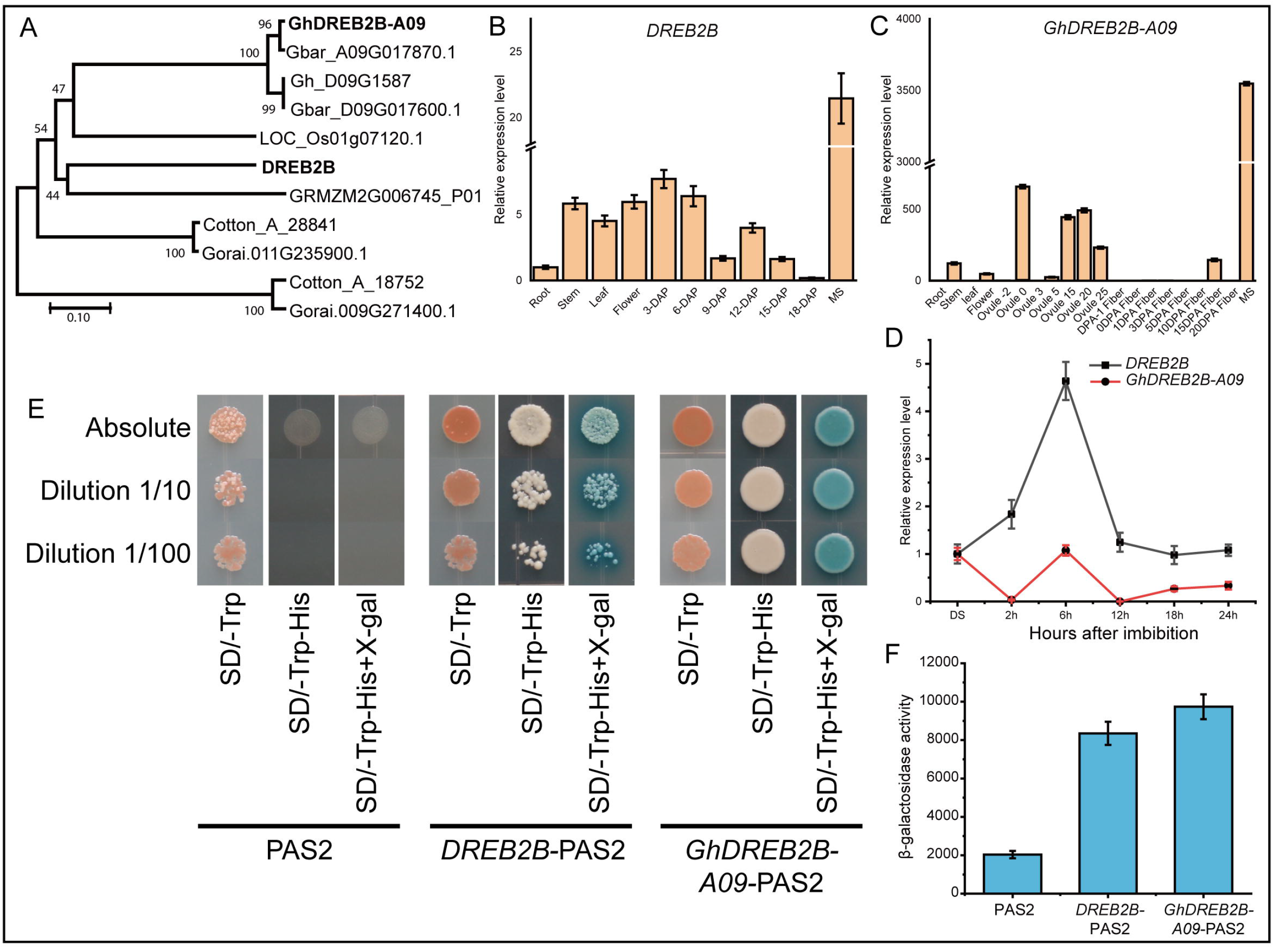
Expression and transcription activity analysis of DREB2B and GhDREB2B-A09 transcription factor. (A) Phylogenetic analysis of DREB2B and its homologous proteins. Phylogenetic tree of DREB2B homolog transcription factors in *G. arboreum, G. hirsutum, G. barbadense, G. raimondii, Z. may*, and *O. sativa*. The neighbor-joining tree based on multiple sequences alignment was built to show the relationships among DREB2B proteins from seven other plant species. (B, C) Determination of mRNA level of *DREB2B* and *GhDREB2B-A09* in different developmental tissues of *Arabidopsis* and cotton respectively. (D) Determination of mRNA level of *DREB2B* and *GhDREB2B-A09* in imbibed seeds of *Arabidopsis* and cotton collected at different time intervals. (B-D) Error bars represent the standard deviations of three independent experiments. (E) Transcriptional activity of DREB2B and GhDREB2B-A09 was determined by a yeast assay approach. 10 and 100 times diluted culture with absolute culture was spread on three different types of plates. The growth of yeast cells on SD/-Trp, SD/-Trp-His, and SD/-Trp-His+ β-galactosidase medium was analyzed. pAS2 (an empty vector) was used as a negative control. (F) the β-galactosidase activity of *pAS2-DREB2B* and *pAS2-GhDREB2B-A09*. β-galactosidase activity from yeast culture of pAS2-*DREB2B* and pAS2-*DREB2B-A09* was calculated in Miller units represented at y-axis and the error bars denoted the standard deviations of three independent experiments.

Furthermore, the expression patterns of *DREB2B* and *GhDREB2B-A09* were examined by qRT-PCR in different developmental tissues and imbibed seeds. The results showed that *DREB2B* and *GhDREB2B-A09* were preferentially expressed in the mature seed as compared to other tissues (Fig. 3B, C). In imbibed seeds, the transcript level of *DREB2B* was up-regulated gradually and reach a peak until 6 h after imbibition, after that it was down-regulated to the level of dry seed; while the transcript level of *GhDREB2B-A09* was equally in dry seeds and at 6 h imbibition, and decreased clearly in other imbibition time points detected (Fig. 3D). These results suggested that *DREB2B* and *GhDREB2B-A09* may share some conserved role partially in seed germination and vigor, the difference in expression profiles of both also indicated potential functional differentiation of DREB2B in Arabidopsis and cotton corresponding with their different amino acid sequences.

### Transcriptional activity of DREB2B in Arabidopsis and cotton

Both DREB2B and GhDREB2B-A09 belongs to the AP2 domain transcription factor family. Therefore, their transcriptional activity was tested by a yeast assay system as described in “Materials and Methods” section. Both transcription factors showed obvious transcriptional activity than the control, and the GhDREB2B-A09 acquired higher transcriptional activity than DREB2B (Fig. 3E). Furthermore, the transcriptional activity was also confirmed by β-galactosidase assay and the results agreed with the yeast assay (Fig. 3F). Collectively, these results illustrated and confirmed the transcriptional activity of both transcription factors from two different species *(Arabidopsis* and *G. hirsutum)*.

### DREB2Bs localization in different sub-cellular or tissue level

To further investigate the mechanisms of DREB2Bs, the transient expression of DREB2B and GhDREB2B-A09 at the sub-cellular level was determined in epidermal cells of tobacco leaves and fusion proteins green fluorescent signals were detected in the membrane and cytoplasm (Fig. 4A).

**Fig. 4:**
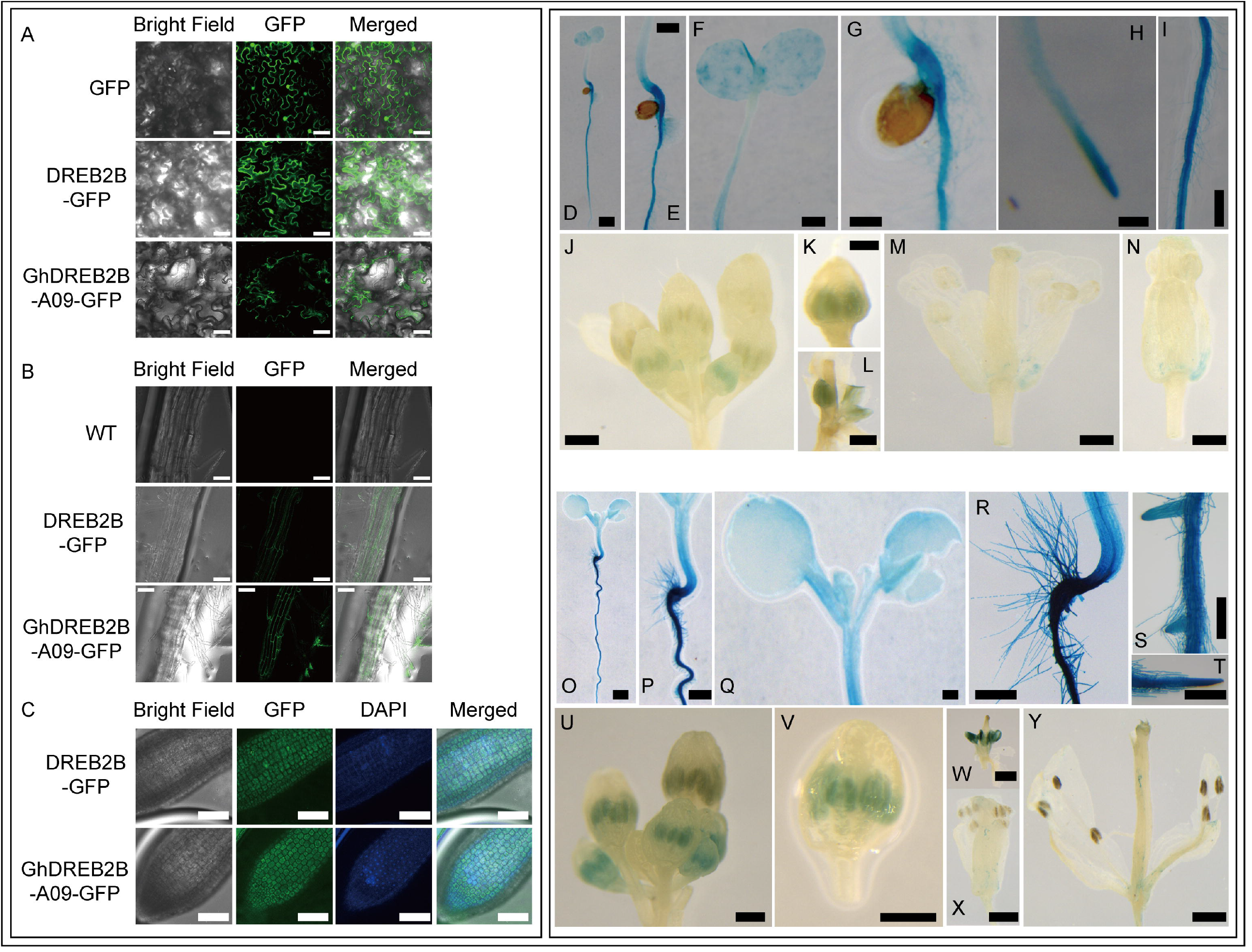
Sub-cellular and tissue level localization of *DREB2B* and *GhDREB2B-A09*. (A) GFP-DREB2B and GFP-GhDREB2B-A09 were transiently expressed in tobacco epidermal cells and green fluorescence (GFP) was observed with a confocal microscope. The images were presented bright field, fluorescence, and merge of bright field and fluorescence. (B, C) GFP-DREB2B and GFP-GhDREB2B-A09 with GFP was transformed in Col and stable expression of green fluorescence (GFP) was observed endogenously in the root and radical cells after 24 hours of imbibed of transgenic plants at T4 generation. The images were presented bright field, fluorescence, and merge of bright field and fluorescence. DAPI, 4,6-diamidino-2-phenylindole. (D-Y) *DREB2B* and *GhDREB2B-A09* promoter activity were determined by histochemical GUS staining. (D, O) Ten-day-old Seedlings, (E, P) shoot and root, (F, Q) Leaf, (G, R) Shoot and root initiation point, (I, S) Root, (H, T) Root tip, (J-L, U-W) anther, (N, X) petals, (M, Y) stigma. (A, B) Bars, 50 µm. (C-Y) Bars, 500µm.

To verify this type of localization in the cytoplasm instead of the nucleus, the 15-days old seedlings root cells of over-expressed *DREB2B-GFP* and *GhDREB2B-A09-GFP* transgenic plants at T4 generation were examined again. The green fluorescent signals in young seedlings’ root cells were also detected in the cytoplasm and membrane (Fig. 4B). Furthermore, the cytoplasm localization was also confirmed by detecting green fluorescent signals with DAPI staining in the radical cells of transgenic plant seeds after 24 h imbibition (Fig. 4C). In combination, these results proved and authenticated the multiple sub-cellular localizations (e.g. cytoplasm, membrane) of DREB2B and GhDREB2B-A09.

Further, 2KB promoter sequences of *DREB2B* and *GhDREB2B-A09* from the translational start site (TSS) was used for promoter *cis*-elements analysis which showed that both genes contained hormones (MeJA, salicylic acid, gibberellin, auxin, and abscisic acid), light, defense and stressed, plant growth and development (endosperm and meristem expression) responsive and MYBHv1 binding sites related *cis*-elements in their promoter region (Supplementary Fig. S3C). Then, promoter activities of *DREB2B* and *GhDREB2B-A09* at T4 generation were investigated in all vegetative and reproductive stages by histochemical GUS staining. *DREB2B* and *GhDREB2B-A09* showed strong promoter activity in almost all vegetative and reproductive tissues such as in young seedlings, shoot, root, leaf, shoot and root initiation point, root tip, root hair, secondary root, anther, petals, and stigma (Fig. 4D-Y) respectively, indicating the similar and constitutive function of both genes in seedling and reproductive organs development of the plant.

### Over-expression of DREB2B and GhDREB2B-A09 reduce seed vigor

To deeply investigate the role of *DREB2B* in seed germination and vigor of Arabidopsis and cotton, *DREB2B* and *GhDREB2B-A09* over-expression lines *(35S:DREB2B* and *35S:GhDREB2B-A09)* were generated. qRT-PCR results showed a significantly higher expression of *DREB2B* and *GhDREB2B-A09* in over-expression lines of both genes, respectively (Supplementary Fig. S4A, B). Moreover, the upregulated expression of *ABI3* in *35S:DREB2B* lines elucidating that *ABI3* is regulated by *DREB2B* at least partially to involve the regulation of seed germination and vigor.

Further, germination speed of *35S:DREB2B* and *35S:GhDREB2B-A09* lines, mutant and WT were tested. The germination of *dreb2b* was earlier than WT, while *35S:DREB2B* and *35S:GhDREB2B-A09* lines showed delayed germination compared to WT at 96 and 120 h imbibition. The *dreb2b* mutant seeds rapidly started to germinate and the germination rate reached over 60% and 80% at 96 h and 120 h imbibition respectively. WT seeds began to germinate later and reached about 10% and 65% germination rate at 96 h and 120 h respectively, while the *35S:DREB2B* and *35S:GhDREB2B-A09* lines began to germinate at 96 h and reached about 35% germination rates at 120 h. Moreover, two *DREB2B* complementary lines exhibited a similar seed germination speed to that of WT. The above results validated the conservative and negative relationship between seed germination and *DREB2B* expression in Arabidopsis and cotton (Fig. 5A, B).

**Fig. 5:**
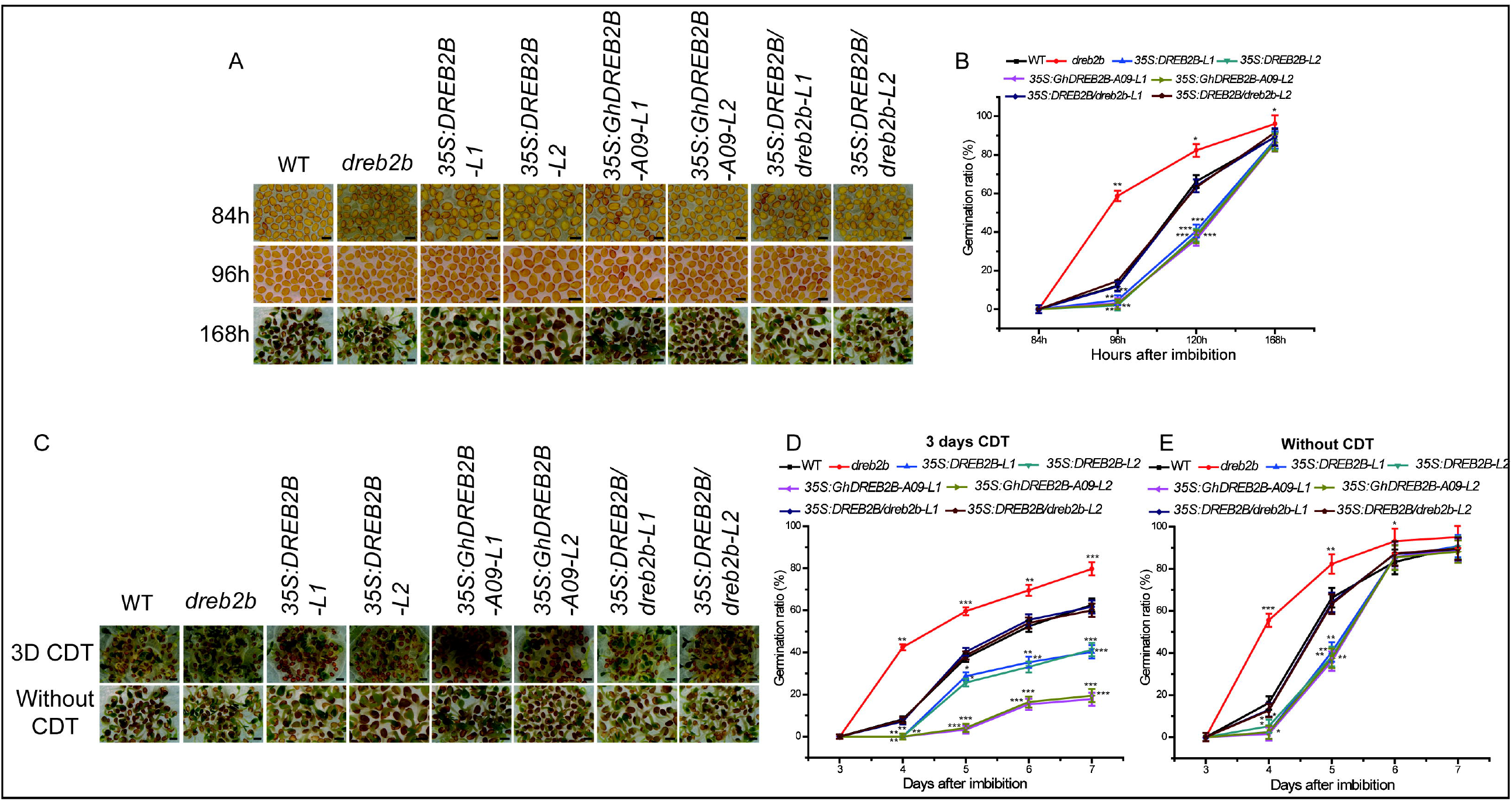
Seed germination and vigor phenotype of WT, *dreb2b* mutant, and complementary lines (*35S:DREB2B/dreb2b*), *35S:DREB2B*, and *35S:GhDREB2B-A09* lines. (A) Germinating seeds microscopic images of WT, *dreb2b* mutant and complementary line, *35S:DREB2B* and *35S:GhDREB2B-A09* lines were taken at 84, 96 and 168 hours after imbibition followed stratification. For each genotype, 80-100 seeds were used. Bars, 500µm. (B) Seed germination after different hour intervals were observed and calculated for WT, *dreb2b* mutant, complementary lines, *35S:DREB2B*, and *35S:GhDREB2B-A09* lines from fresh seeds harvested 2 weeks before the experiment. (C) Seed germination images of WT, *dreb2b* mutant, and complementary lines, *35S:DREB2B* and *35S:GhDREB2B-A09* lines treated with or without CDT treatment. Images were taken 7 days after imbibition with stereo-microscope. Bars, 500µm. (D, E) WT, *dreb2b* mutant and complementary lines, *35S:DREB2B* and *35S:GhDREB2B-A09* lines seeds germination ratio treated with 3 days and without CDT was calculated at different time points. (B, D, and E) Experiments were performed in three replicates and germination ratio was measured from averages of three replicates with about 100 seeds per replicate from independent lines of each genotype. Error bars indicate SD based on three biological replicates. Asterisks over the bars indicate a significance level among means determined by the student’s T-test (*P□ 0.05, **P□ 0.01, ***P□ 0.001).

In turn, CDT assay also was used to confirm the seed vigor and viability of *35S:DREB2B* and *35S:GhDREB2B-A09* lines. After 3-days CDT treatment, seeds of all the genotypes exhibited some delay in germination compared with mock; while *dreb2b* mutant seeds showed a relatively lower sensitivity, and *35S:DREB2B* and *35S:GhDREB2B-A09* lines showed higher sensitivity, as compared with WT (Fig. 5C, D). The germination ratio was 40% and 17-19% for *35S:DREB2B* and *35S:GhDREB2B-A09* lines respectively, whereas WT and *dreb2b* showed 60% and 81% germination at 7-days imbibition respectively. A tetrazolium assay after 3-day CDT treatment showed that viable seeds in *35S:DREB2B* and *35S:GhDREB2B-A09* lines were decreased to a lower ratio 6-13%; while *dreb2b* mutant displayed more viable seeds (42%) compared to WT (19%) and complementary lines (17-20%) (Supplementary Fig. S4B, C).

### DREB2B interacts with RCD1conservatively in Arabidopsis and cotton

The above results showed that DREB2B functions dependent and independent on ABI3 to mediate the seed vigor, which forwards us to unravel some interacting factors of DREB2B to understand a more detailed pathway involved by DREB2B in seed vigor. Online prediction analysis (eFP Browser and ccNET) indicated that DREB2B protein might interact with RCD1 (RADICAL-INDUCED CELL DEATH 1) and SRO1 (SIMILAR TO RCD-ONE 1) in Arabidopsis and cotton. To confirm that, five homologous RCDs *(GhRCD1L-A05, GhRCD1L-D05, GhRCD1L-A08, GhRCD1L-A12*, and *GhRCD1L-D12)* (Supplementary Table S2) were identified in cotton and the alignment of the amino acid of total seven RCD1 family proteins showed the moderate identity among them; then a phylogenetic tree was constructed and depicted that GhRCD1s in chromosomes A12, D12 and A08 are the more closer homology with *Arabidopsis* RCD1 and SRO1 (Supplementary Fig. S5A, B). Further analysis indicated that all RCD1s from *Arabidopsis* and *G. hirsutum* contain WWE domain and other two domains (PARP and RST), which may contribute to the conserved and physical interaction (Supplementary Fig. S5C) between DREB2B and RCD1 family genes in the plant.

Then, the interaction of DREB2B proteins with RCD1 and SRO1 in *Arabidopsis* and cotton were subsequently analyzed in *planta* using BiFC. The interaction of Arabidopsis DREB2B with RCD1 was observed in the membrane and cytoplasm while with SRO1 in the nucleus (Fig. 6A). However, the interactions of GhDREB2B-A09 with five GhRCD1Ls proteins (GhRCD1L-A05, GhRCD1L-D05, GhRCD1L-A08, GhRCD1L-A12, and GhRCD1L-D12) were observed in the membrane and cytoplasm (Fig. 6B). These results were in accordance with the previous study that *RCD1* and *SRO1* interacted with several transcription factors belonging to the *DREB* transcription factor family including *DREB2B* and other transcription factor families (Jaspers *et al*., 2009; Wu *et al*., 2018). Meantime, the localization difference between the interaction factor analysis of DREB2B from Arabidopsis and cotton indicated some specificity for the underlying mechanisms associated with DREB2B in seed vigor regulation of different plant species.

**Fig. 6:**
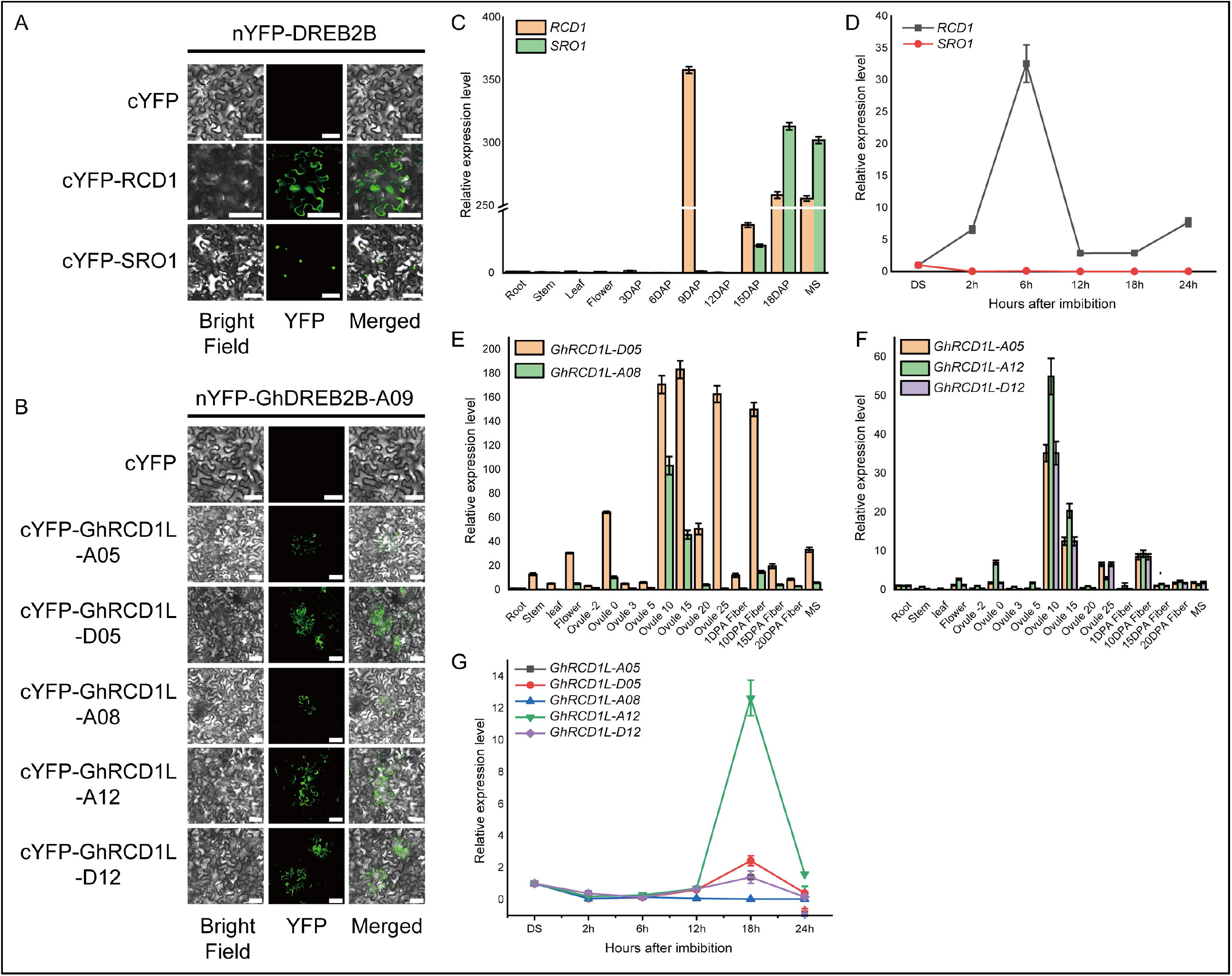
Interaction between DREB2B and the member of RCD1 family proteins in cotton and *Arabidopsis* and the expression profiles of *RCD1* family genes in cotton and Arabidopsis. (A) Bimolecular fluorescence complementation (BiFC) for interactions between RCD1, SRO1 and DREB2B. For BiFC assays, *Nicotiana benthamiana* leaves were co-agro-infiltrated with constructs expressing *DREB2B* fused to the N- and *RCD1*or *SRO1* fused to the C-terminus half of YFP. *RCD1* showed strong interaction with *DREB2B* in the cytoplasm while *SRO1* in the nucleus. Bars, 50µm. (B) BiFC assay for interactions between *GhRCD1L* and *GhDREB2B-A09* was performed as described before. *GhRCD1L-A05, GhRCD1L-D05, GhRCD1L-A08, GhRCD1L-A12*, and *GhRCD1L-D12* showed strong interaction with *GhDREB2B-A09* in the cytoplasm. The reconstructed YFP fluorescence was recorded 72h post agroinfiltration by confocal microscopy. Bars, 50µm. (C, E and F) mRNA level of *RCD1, SRO1*, and *GhRCD1L* in different developmental tissues of *Arabidopsis* and cotton. (D, G) mRNA level of *RCD1, SRO1*, and *GhRCD1L* in imbibed seeds of *Arabidopsis* and cotton collected at different time intervals. (C-G) Error bars represent the standard deviations of three independent experiments. *ACTIN2* and *GhHis3* genes were used as an internal control for *Arabidopsis* and cotton respectively.

### RCD1 and SRO1 negatively regulate seed vigor

To investigate the roles of *RCD1* and *SRO1*, transcript levels of them were determined in different developmental tissues and imbibed seeds of *Arabidopsis* and *G. hirsutum. RCD1* exhibited very higher transcript levels (no less than 200 folds higher compared to other tissues) in 9, 15, 18 DAP siliques and mature seeds. *SRO1* transcript level was gradually up-regulated during the late phase of seed development at 15 and 18 DAP siliques and mature seeds similar to *RCD1* (Fig. 6C). The up-regulated expression of *RCD1* and *SRO1* during seed maturation indicated that both genes have specified roles during seed maturation. In addition, *RCD1* exhibited almost the same pattern as that of *DREB2B* in imbibed seeds (Fig. 6D and 3D), suggesting that *RCD1* might function in seed germination similar to *DREB2B*. However, the *SRO1* mRNA was down-regulated in imbibed seeds at all-time points (Fig. 6D).

In cotton, *GhRCD1L-A05* and *GhRCD1L-D05* showed constitute expression and higher levels in later developmental ovules; *GhRCD1L-A08, GhRCD1L-A12*, and *GhRCD1L-D12* showed specific expression in the ovules (Fig. 6E, F). While in imbibed seeds, the only expression of *GhRCD1L-A12* was up-regulated clearly at 18 h imbibition (Fig. 6G). The expression results indicated that *RCD1* family genes may play redundant and specific roles in seed related traits.

To confirm the roles of *RCD1* and *SRO1* in seed germination and vigor, germination speed of *rcd1-3, sro1-1*, and *sro1-2* in freshly harvested seeds was examined at different hour intervals during imbibition after stratification. *rcd1-3, sro1-1*, and *sro1-2* germination were earlier and completed the 50-60% germination in 96 h after stratification similar to *dreb2b* while only 10% seed germination was observed in WT (Fig. 7A, B). More than 80% seed germination of *dreb2b, rcd1-3, sro1-1*, and *sro1-2* mutants were observed, whereas, WT showed less than 60% germination after 120 h imbibition. The similar seed germination phenotype of *dreb2b, rcd1-3, sro1-1*, and *sro1-2* mutants demonstrated that DREB2B and its interacting proteins (RCD1 and SRO1) may function through the same pathway to involve seed germination.

**Fig. 7:**
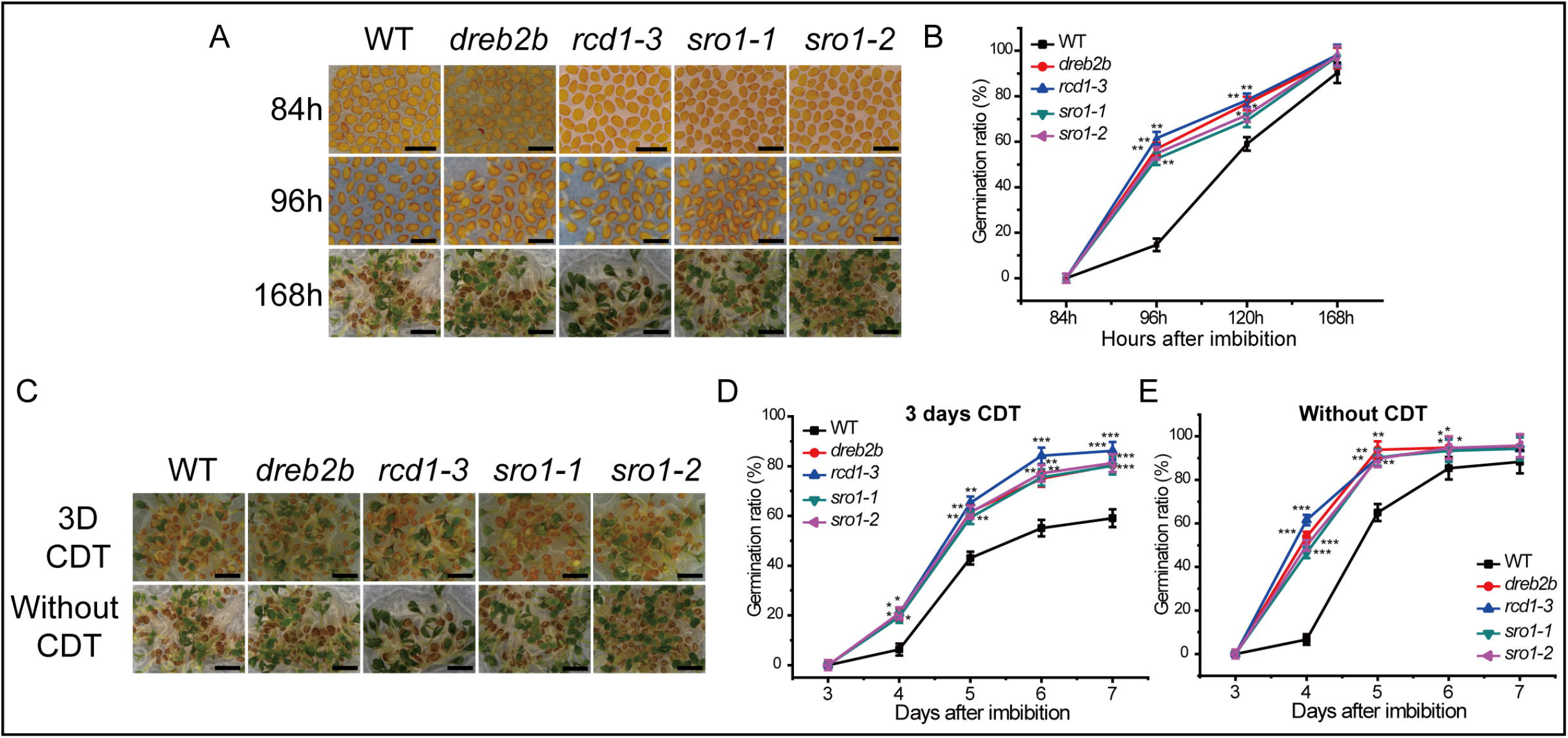
Phenotypic analysis of seed vigor of WT, *dreb2b, rcd1-3, sro1-1* and *sro1-*2 (A) Seeds microscopic images of WT, *dreb2b, rcd1-3, sro1-1* and *sro1-*2 showing germination speed were taken at 84, 96 and 168 hours after imbibition followed stratification. For each genotype, 80-100 seeds were used. Bars, 1mm. (B) Germination ratio after different hour intervals was observed and calculated for WT, *dreb2b, rcd1-3, sro1-1*, and *sro1-*2 from fresh seeds harvested 2 weeks before the experiment. (C) Images of WT, *dreb2b, rcd1-3, sro1-1*, and *sro1-*2 seeds treated with or without CDT treatment were taken at 7 days after imbibition with stereo-microscope. For each genotype, 80-100 seeds were used. Bars, 1mm. (D, E) WT, *dreb2b, rcd1-3, sro1-1* and *sro1-*2 seed germination ratio treated with 3 days and without CDT was calculated with different day’s interval. (B, D, and E) For each genotype, three replicates with about 100 seeds per replicate were used and error bars represented averages ± s.d. of three independent batches of seeds for each genotype. Asterisks over the bars indicate a significance level among means determined by the student’s T-test (*P□ 0.05, **P□ 0.01, ***P□ 0.001).

Furthermore, *rcd1-3, sro1-1*, and *sro1-2* mutants showed less sensitivity to CDT compared to WT similar to *dreb2b*, and germination ratio of treated seeds was 59, 80, 90, 80, and 81% for WT, *dreb2b, rcd1-3, sro1-1*, and *sro1-2* respectively after 7-days (Fig. 7C, D) imbibition; all these lines also showed similar seed germination pattern with above without CDT treatment (Fig. 7E, B). These results confirmed that DREB2B together with RCD1 and SRO1 is playing a negative role in the regulation of seed vigor and germination.

Here *rcd1-3, sro1-1*, and *sro1-2* mutants showed similar seed vigor and germination phenotype to *dreb2b* and confirmed that *DREB2B* works together with *RCD1* and *SRO1* to negatively involve seed vigor and germination regulation. Hence, to further confirm the correlation between DREB2B and its interacting proteins (RCD1 and SRO1), the expression of RCD1 and SRO1 in *dreb2b, abi3-16* and *dreb2b/abi3-16* mutant’s dry seeds while DREB2B and ABI3 expression in *rcd1-3*, and *sro1-1* mutant’s dry seeds was detected (Supplementary Fig. S6A, B). The expression analysis exhibited the 0.7 to 0.8 fold down-regulated mRNA level of *RCD1* and *SRO1* in *dreb2b* and *abi3-16* mutant’s dry seeds while their expression was almost completely down-regulated in *dreb2b/abi3-16* double mutant dry seeds. However, the transcript level of *DREB2B* and *ABI3* in *rcd1-3*, and *sro1-1* was also down-regulated confirmed that both interacting proteins (DREB2B, RCD1, and SRO1) positively regulate each other and work together in the regulation of seed vigor and germination negatively. Besides, the up-regulated expression of RCD1 and SRO1 in response to ABA treatments showed that similar to DREB2B its interacting proteins also induced by ABA and use ABA-mediated pathway for seed vigor and germination regulation negatively (Supplementary Fig. 6C).

## Discussion

### DREB2B plays a negative role in seed germination and vigor

EREBP, a super-family of plant-specific transcription factors, contain a highly conserved AP2 DNA-binding domain (Okamuro *et al*., 1997; Weigel, 1995), and were classified into subfamilies DREB, ERF, AP2, RAV, and others based on the number of repetitions and the sequence of the AP2 domain in *Arabidopsis* (Sakuma *et al*., 2002). The DREB proteins namely, DREB1 and DREB2 can induce a set of abiotic stress-related genes (Lata and Prasad, 2011). Eight DREB2-type proteins were identified in Arabidopsis (Nakashima *et al*., 2000; Sakuma *et al*., 2002), among which *DREB2B* confers various abiotic stress-tolerance in many species (Marcos-Filho, 2015; Pettigrew and Dowd, 2011). Here, a loss of function *dreb2b* mutant was identified and it exhibited significantly longer seed longevity and quicker germination compared to the WT (Fig. 1A, B, C). Additionally, *dreb2b* mutant showed higher seed germination and more viable seeds than WT after artificial aging treatment (Fig. 1D-H). These results suggested that the abundance of BREB2B protein in seeds appears to be negatively correlated with seed germination and longevity.

### DREB2B is involved in ABA-dependent pathway partially

ABA is a type of very vital hormone in seed development as well as seed vigor (Dekkers *et al*., 2016). In our study, it is observed that the absence of DREB2B protein leads to decreased inhibition of seed germination in response to exogenous ABA, suggesting that *DREB2B* might function in an ABA-dependent pathway to direct seed germination and vigor. Moreover, the significantly reduced expression of *ABI3* and higher expression of *CYP707A1* in *dreb2b* supported that both ABA synthesis and signaling play downstream of DREB2B at least partially (Fig. 2B, Supplementary Fig. S2B). Collectively, the reduced ABA sensitivity of *dreb2b*, induced expression of *DREB2B* in response to ABA treatments, and altered expression of ABA pathway genes all suggested that DREB2B exert roles involved in the ABA pathway to negatively manage seed germination and vigor (Fig. 2A, B).

### DREB2B functions synergistically with ABI3 in seed vigor regulation

ABI3 is a well-known positive regulator for seed vigor and seed dormancy (Arad *et al*., 2002; Ooms *et al*., 1993), supporting the results in this study (Fig. 2C-H). The down-regulation of *ABI3* in *dreb2b* and unchanged expression of *DREB2B* in *abi3-16* indicated that *DREB2B* might function upstream of *ABI3* partially (Fig. 2B, Supplementary Fig. S2B). Moreover, *dreb2b/abi3-16* double mutant exhibited better seed germination and vigor than single mutants (Fig. 2D, F, G) which indicated that the DREB2B and ABI3 show synergistic roles in genetic in the seed germination speed and seed vigor through different pathways. Combining with the expression of *DREB2B* or *ABI3* in *abi3-16* or *dreb2b* mutants respectively, all these indicated that seed germination after stratification and seed germination after CDT treatment may not be regulated by the same pathways, and in these different processes, DREB2B and ABI3 play roles with an additive model in part as well as through some different downstream pathways.

### Conservative roles of DREB2B in seed vigor of Arabidopsis and cotton

To ask the function of *DREB2B* in cotton seed vigor, *GhDREB2B-A09* was identified as the homologous gene of *DREB2B* from *G. hirsutum* with the conserved AP domains (Supplementary Fig. S3A, B). The expression analysis indicated that the higher expression of *DREB2B* and *GhDREB2B-A09* were detected in mature seed compared to other tissues of plant development. Moreover, in imbibed seeds, the expression of *DREB2B* and *GhDREB2B-A09* were up-regulated quickly. The similar higher transcript level in mature seed and during imbibition suggested that *DREB2B* and *GhDREB2B-A09* are playing similar promising roles in seed germination and vigor (Fig. 3B-D). Besides, transcriptional activity through the yeast system and β-galactosidase assay illustrated that DREB2B and GhDREB2B-A09 are functional transcription activation factors (Fig. 3E, F).

Next, *35S:DREB2B* and *35S:GhDREB2B-A09* lines showed significantly slow germination speed (Fig. 5A, B) and drastically reduced seed vigor and viability after CDT treatment compared to WT, complementary lines, and *dreb2b* (Fig. 5C-E, Supplementary Fig. S4B, C). All these supported and confirmed that DREB2B is negatively involved in seed vigor and germination in *Arabidopsis* and cotton.

### How is the biochemical and molecular channels of DREB2B in seed germination and vigor regulation?

DREB2B is involved in multiple abiotic stresses tolerance and seed development. However, its detailed biochemical and molecular mechanism is still ambiguous. Firstly, we tested the protein sub-cellular localization. Transient and stable expression analysis of protein localization of DREB2B and GhDREB2B-A09 authenticated that both DREB2B transcription factor proteins are exclusively localized in cytoplasm and membrane, indicating the different mechanisms of DREB2B from common transcription factors in the nucleus, which also provide some new and conceivable molecular mechanisms of transcription factors. Both of the promoters-GUS analysis of *DREB2B* and *GhDREB2B-A09* also showed their similar and constitute expression in plant development (Fig. 4D-Y), supporting the multiple and conservative roles of DREB2B in Arabidopsis and cotton. Second, the interaction factors of DREB2B and GhDREB2B-A09, RCD1s were explored and identified by protein interaction software (eFP Browser and ccNET) and supported the previous study (Wu *et al*., 2018). Further, BiFC experiments proved the direct interaction of DREB2B and GhDREB2B-A09 with RCD1s in membrane, cytoplasm, or nucleus, which indicated the potentially different molecular mechanisms of DREB2B associated with different interacting factors. But it remains to be determined the putative interaction specificity between two proteins and the biological significance of their interactions. RCD1 belongs to the (ADP-ribosyl) transferase domain containing subfamily with a WWE protein-protein interaction domain and acts as an integrative node within different hormonal signaling (e.g. ABA, Ethylene, JA) and in several stresses responses (e.g. drought, cold) (Ahlfors *et al*., 2004). In addition, the interaction of RCD1 with other proteins highlighted its different functions. For example, RCD1 interacts with the cytoplasmic tail of Salt Overly Sensitive1 (SOS1) which elucidated the cross-talk between the ion-homeostasis and oxidative-stress detoxification pathways involved in plant salt tolerance (Katiyar-Agarwal *et al*., 2006). In another study, the interaction between glutathione peroxidase3 and RCD1 was identified which hypothesized that RCD1 could be a plant equivalent of the yeast redox-regulated transcription factor Yap1 (Miao *et al*., 2006). Moreover, previous screening of RCD1 interaction factor by yeast two-hybrid identified several transcription factors such as DREB2A, DREB2B (Belles-Boix *et al*., 2000), suggesting that RCD1 might affect the function or activity of transcription factors to regulate downstream genes, which also support the interaction between RCD1 and DREB2B in seed development regulation. Collectively, RCD1 is a multifunctional factor interacting with various proteins, which make it like an adaptor or post-translational factor involved in the different complex and physiological pathway, but the detailed mechanisms are unclear.

To confirm the physiological roles of RCD1 genes in seed germination and vigor, the expressions of *RCD1, SRO1* were investigated in different developmental tissues and imbibed seeds of *Arabidopsis* and *G. hirsutum* by qRT-PCR (Fig. 6C-G). The higher transcript of *RCD1* and S*RO1* during seed maturation and imbibition compared to other tissues indicated their potential roles during seed development and germination. The similar expression pattern of *RCD1* family genes and *DREB2B* in developed and imbibed seeds suggested the underlying interaction between them in seed development and germination. Furthermore, in line with expectations, *rcd1-3, sro1-1*, and *sro1-2* mutants exhibited a quicker germination rate and better seed vigor same as *dreb2b* (Fig. 7A-E). Additionally, the down-regulated expression of *RCD1* and *SRO1* in *dreb2b* and *DREB2B* in *rcd1-3* and *sro1-1* further proved their interaction; up-regulated expression of *RCD1* and *SRO1* in response to ABA similar to DREB2B evidenced the same pathway involved by them (Supplementary Fig. S6A-C). From these results, it is concluded that DREB2B and its interacting proteins (RCD1 and SRO1) have a positive correlation with each other and form complex to regulate seed germination and vigor negatively together via an ABA-mediated pathway.

### Hypothetical model associated with DREB2B in seed vigor/longevity regulation

In a word, we hypothesized a model for DREB2B in seed vigor regulation here (Fig. 8). ABI3 is an important positive regulator for seed vigor through downstream heat shock factors such as HsfA9, which is controlled by DREB2B partially. Furthermore, the complex comprising DREB2B and RCD1s also plays crucial roles synergistically with ABI3 through an unidentified downstream factor or pathway in seed vigor, which functions downstream of ABA same as ABI3. To exploring the downstream factors by RNA-Seq, ChIP-Seq et al. approaches would be very important in the detailed mechanisms involved by DREB2B in seed vigor regulation. A previous study showed that ROS (reactive oxygen species) change the profile and function of the RCD1 protein. RCD1 interacts and suppresses the activity of the transcription factors ANAC013 and ANAC017, which mediate a ROS-related retrograde signal originating from mitochondrial complex III, providing a feedback control on its function (Shapiguzov *et al*., 2019). These indicated that RCD1 can modify interacting factors positively or negatively with several different approaches to mediate downstream pathways. Thus, to reveal the underlying mechanism within the interaction between DREB2B and RCD1 would provide more interesting and novel clues for the roles of them. DREB2B showing a non-nuclear localization indicated its special mechanism. More intensive research such as determination of the structure, transcription activity, and protein localization of DREB2B before and after interaction with RCD1 family proteins would be interesting and helpful for the detailed mechanism of DREB2B and RCD1 in seed germination and vigor. The down-regulated mRNA level of *RCD1* and *SRO1* in *snl1snl2-1* double mutant like *DREB2B* suggested that *SNL1/2* functions upstream of *DREB2B, RCD1*, and *SRO1* in the regulation of seed germination and vigor through an indirect interaction (Supplementary Fig. S6D), which shed some light on the regulation between epigenetic and DREB2B-RCD1 complex.

**Fig. 8:**
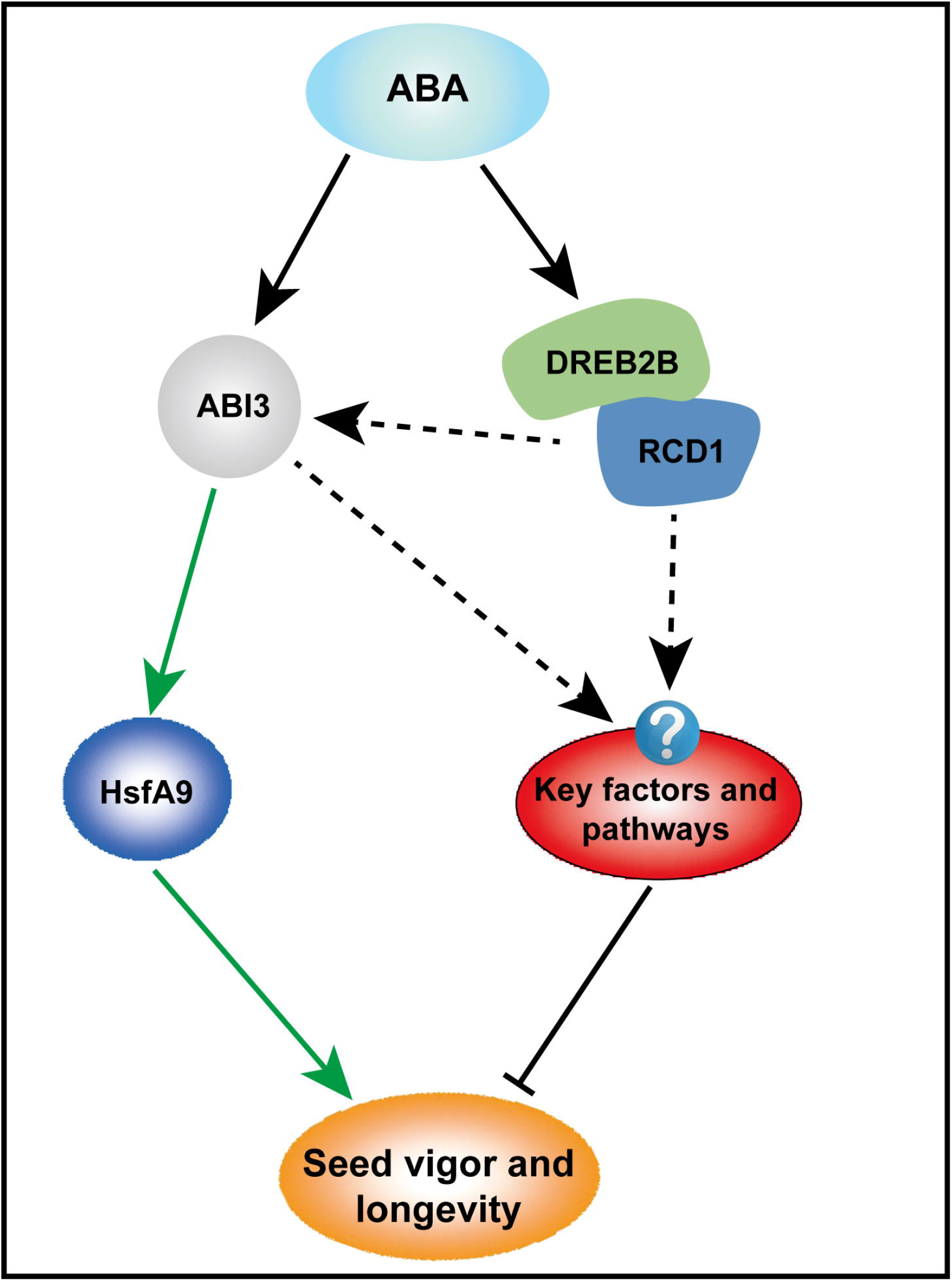
Model of DREB2B function in seed vigor dependent and independent on ABI3 pathways. Previously identified seed vigor regulation pathway illustrated that ABI3 induced the expression of downstream transcription factor HsfA9 to improve seed vigor. Comparatively, in our study, DREB2B interacts and form a complex with RCD1s, meanwhile it partially induce the expression of *ABI3* and function synergistically with ABI3 in seed vigor regulation by activating some unfamiliar factors and pathways. In this model, ABI3 is involved in two antagonistic pathways, one is dependent on HsfA9 positively and another is dependent on an unclear pathway negatively, but the correlation and underlying mechanisms between these two pathways are blank. Anyway, both ABI3 and DREB2B associated pathways are activated by ABA in seed vigor regulation.

## Acknowledgement

We thank Xin Li (ICR, CAAS) for the technical support in using confocal microscopy. This work was supported by grants from the National Natural Science Foundation (31690093), the Creative Research Groups of China (31621005), and Zhengzhou University (32410196).

## Author Contribution

ZW and FL conceived and designed the study. ZW and FA performed research; YL, LG, and Z Y provided reagents and materials; ZW and FA drafted the manuscript. All authors revised and approved the manuscript.

## Data availability statement

All data supporting the findings of this study are available within the paper and within its supplementary materials published online.

## Supporting Information

Supplementary data are available at *JXB* online.

**Table S1:** Information related mutants that are used in the study.

**Table S2:** Primers for mutants screening, qPCR, and amplification of coding regions and promoters of genes used in the study.

**Fig. S1:** Identification of *dreb2b* null mutant and its seed germination and vigor phenotype verification.

**Fig. S2:** *DREB2B* is induced by ABA to involve in seed vigor regulation.

**Fig. S3:** Protein and promoter sequences analysis of DREB2B and GhDREB2B-A09.

**Fig. S4:** Expression of *DREB2B* and *GhDREB2B-A09* in over-expression lines and determination of seed germination and vigor of WT, *dreb2b* mutant, and complementary lines, *35S:DREB2B* and *35S:GhDREB2B-A09* lines.

**Fig. S5:** Protein sequence analysis of RCD1s in Arabidopsis and cotton.

**Fig. S6:** Expression analysis of *DREB2B, ABI3, RCD1*, and *SRO1* in different lines and response to ABA.

## References

Abdelmagid AS, Osman AM. 1975. Influence of Storage Period and Temperature on Viability and Chemical Composition of Cotton Seeds. Annals of Botany 39, 237–248.

Ahlfors R, Lång S, Overmyer K, Jaspers P, Brosché M, Tauriainen A, Kollist H, Tuominen H, Belles-Boix E, Piippo M, Inzé D, Palva ET, Kangasjärvi J. 2004. Arabidopsis RADICAL-INDUCED CELL DEATH1 Belongs to the WWE Protein–Protein Interaction Domain Protein Family and Modulates Abscisic Acid, Ethylene, and Methyl Jasmonate Responses. The Plant cell 16, 1925.

Ali F, Qanmber G, Wei Z, Yu D, Li Yh, Gan L, Li F, Wang Z. 2020. Genome-wide characterization and expression analysis of geranylgeranyl diphosphate synthase genes in cotton (Gossypium spp.) in plant development and abiotic stresses. BMC Genomics 21, 561.

Angelovici R, Galili G, Fernie AR, Fait A. 2010. Seed desiccation: a bridge between maturation and germination. Trends in Plant Science 15, 211–218.

Arad M, Benson DW, Perez-Atayde AR, McKenna WJ, Sparks EA, Kanter RJ, McGarry K, Seidman J, Seidman CE. 2002. Constitutively active AMP kinase mutations cause glycogen storage disease mimicking hypertrophic cardiomyopathy. The Journal of clinical investigation 109, 357–362.

Bailly C, Leymarie J, Lehner A, Rousseau S, Côme D, Corbineau F. 2004. Catalase activity and expression in developing sunflower seeds as related to drying. J Journal of Experimental Botany 55, 475–483.

Bartee SN, Krieg DR. 1974. Cottonseed Density: Associated Physical and Chemical Properties of 10 Cultivars1. Agronomy Journal 66, 433–435.

Belles-Boix E, Babiychuk E, Van Montagu M, Inzé D, Kushnir S. 2000. CEO1, a new protein from Arabidopsis thaliana, protects yeast against oxidative damage11CEO1 was deposited in the EMBL databank with the accession number AJ251578. FEBS Letters 482, 19–24.

Bentsink L, Jowett J, Hanhart CJ, Koornneef M. 2006. Cloning of DOG1, a quantitative trait locus controlling seed dormancy in Arabidopsis. Proceeding of National Academy of Science U S A 103, 17042–17047.

Cantoro R, Crocco CD, Benech-Arnold RL, Rodríguez MV. 2013. In vitro binding of Sorghum bicolor transcription factors ABI4 and ABI5 to a conserved region of a GA 2-OXIDASE promoter: possible role of this interaction in the expression of seed dormancy. Journal of Experimental Botany 64, 5721–5735.

Chen HH, Chu P, Zhou YL, Ding Y, Li Y, Liu J, Jiang LW, Huang SZ. 2016. Ectopic expression of NnPER1, a Nelumbo nucifera 1-cysteine peroxiredoxin antioxidant, enhances seed longevity and stress tolerance in Arabidopsis. Plant Journal 88, 608–619.

Clerkx EJM, Blankestijn-De Vries H, Ruys GJ, Groot SPC, Koornneef M. 2004. Genetic differences in seed longevity of various Arabidopsis mutants. Physiologia Plantarum 121, 448– 461.

Dekkers BJW, He H, Hanson J, Willems LAJ, Jamar DCL, Cueff G, Rajjou L, Hilhorst HWM, Bentsink L. 2016. The Arabidopsis DELAY OF GERMINATION 1 gene affects ABSCISIC ACID INSENSITIVE 5 (ABI5) expression and genetically interacts with ABI3 during Arabidopsis seed development. The Plant Journal 85, 451–465.

Faiza A, Qanmber G, Yonghui L, Shuya M, Lili L, Zuoren Y, Zhi W, Fuguang L. 2019. Genome-wide identification of Gossypium INDETERMINATE DOMAIN genes and their expression profiles in ovule development and abiotic stress responses. Journal of Cotton Research 2, 3.

Finkelstein R, Reeves W, Ariizumi T, Steber C. 2008. Molecular aspects of seed dormancy. Annual Review of Plant Biology 59, 387–415.

Finkelstein RR, Gampala SS, Rock CD. 2002. Abscisic acid signaling in seeds and seedlings. The Plant Cell 14, S15–S45.

Forman M, Jensen WA. 1965. Respiration and Embryogenesis in Cotton. Plant Physiology 40, 765–769.

Ghaderi-Far F, Khavari F, Sohrabi B. 2012. Lint yield and seed quality response of drip irrigated cotton under various levels of water. International Journal of Plant Production 6.

Hay FR, Valdez R, Lee JS, Sta Cruz PC. 2019. Seed longevity phenotyping: recommendations on research methodology. Journal of Experimental Botany 70, 425–434.

Hu W, Chen M-l, Wenqing Z, Chen B-l, Wang Y-h, Wang S-s, Meng Y-l, Zhou Z-g. 2017. The effects of sowing date on cottonseed properties at different fruiting-branch positions. Journal of Integrative Agriculture 16, 1322–1330.

Jaspers P, Blomster T, Brosché M, Salojärvi J, Ahlfors R, Vainonen JP, Reddy RA, Immink R, Angenent G, Turck F, Overmyer K, Kangasjärvi J. 2009. Unequally redundant RCD1 and SRO1 mediate stress and developmental responses and interact with transcription factors. The Plant Journal 60, 268–279.

Jing-bao L, Zhi-yuan F, Hui-ling X, Yan-min H, Zong-hua L, Liu-jing D, Shang-zhong X, Ji-hua T. 2011. Identification of QTLs for maize seed vigor at three stages of seed maturity using a RIL population. Euphytica 178, 127–135.

K R. 2007. Effect of seed treatments on viability and vigour of cotton seeds (Gossypium hirsutum L.) under ambient storage. Journal of Cotton Research and Development 31, 1–6.

Katiyar-Agarwal S, Zhu J, Kim K, Agarwal M, Fu X, Huang A, Zhu JK. 2006. The plasma membrane Na+/H+ antiporter SOS1 interacts with RCD1 and functions in oxidative stress tolerance in Arabidopsis. Proceeding of National Academy of Science U S A 103, 18816–18821.

Koornneef M, Bentsink L, Hilhorst H. 2002. Seed dormancy and germination. Current opinion in plant biology 5, 33–36.

Kotak S, Vierling E, Bäumlein H, von Koskull-Döring P. 2007. A novel transcriptional cascade regulating expression of heat stress proteins during seed development of Arabidopsis. The Plant cell 19, 182–195.

Kumar S, Stecher G, Tamura K. 2016. MEGA7: Molecular Evolutionary Genetics Analysis Version 7.0 for Bigger Datasets. Molecular Biology and Evolution 33, 1870–1874.

Lata C, Prasad M. 2011. Role of DREB in regulation of abiotic stress response in plants. Journal of experimental botany 62, 4731–4748.

Liu J, Meng Y, Chen J, Lv F, Ma Y, Chen B, Wang Y, Zhou Z, Oosterhuis DM. 2015. Effect of late planting and shading on cotton yield and fiber quality formation. Field Crops Research 183, 1–13.

Liu Q, Kasuga M, Sakuma Y, Abe H, Miura S, Yamaguchi-Shinozaki K, Shinozaki K. 1998. Two transcription factors, DREB1 and DREB2, with an EREBP/AP2 DNA binding domain separate two cellular signal transduction pathways in drought- and low-temperature-responsive gene expression, respectively, in Arabidopsis. The Plant cell 10, 1391–1406.

Liu X, Zhang H, Zhao Y, Feng Z, Li Q, Yang HQ, Luan S, Li J, He ZH. 2013. Auxin controls seed dormancy through stimulation of abscisic acid signaling by inducing ARF-mediated ABI3 activation in Arabidopsis. Proceeding of National Academy of Science U S A 110, 15485–15490.

Lovegrove A, Hooley R. 2000. Gibberellin and abscisic acid signalling in aleurone. Trends in Plant Science 5, 102–110.

Marcos-Filho J. 2015. Seed vigor testing: An overview of the past, present and future perspective. Scientia Agricola 72, 363–374.

Matilla AJ, Carrillo-Barral N, Rodríguez-Gacio MdC. 2015. An Update on the Role of NCED and CYP707A ABA Metabolism Genes in Seed Dormancy Induction and the Response to After-Ripening and Nitrate. Journal of Plant Growth Regulation 34, 274–293.

Miao Y, Lv D, Wang P, Wang X-C, Chen J, Miao C, Song C-P. 2006. An Arabidopsis glutathione peroxidase functions as both a redox transducer and a scavenger in abscisic acid and drought stress responses. The Plant cell 18, 2749–2766.

Nakashima K, Shinwari ZK, Sakuma Y, Seki M, Miura S, Shinozaki K, Yamaguchi-Shinozaki K. 2000. Organization and expression of two Arabidopsis DREB2 genes encoding DRE-binding proteins involved in dehydration- and high-salinity-responsive gene expression. Plant Molecular Biology 42, 657–665.

Nakashima K, Yamaguchi-Shinozaki K. 2010. Promoters and Transcription Factors in Abiotic Stress-Responsive Gene Expression. In: Pareek A, Sopory SK, Bohnert HJ, eds. Abiotic Stress Adaptation in Plants: Physiological, Molecular and Genomic Foundation. Dordrecht: Springer Netherlands, 199–216.

Nambara E, Okamoto M, Tatematsu K, Yano R, Seo M, Kamiya Y. 2010. Abscisic acid and the control of seed dormany and germination. Seed Science Research 20.

Okamuro JK, Caster B, Villarroel R, Van Montagu M, Jofuku KD. 1997. The AP2 domain of APETALA2 defines a large new family of DNA binding proteins in Arabidopsis. Proceeding of National Academy of Science U S A 94, 7076–7081.

Ooms J, Leon-Kloosterziel KM, Bartels D, Koornneef M, Karssen CM. 1993. Acquisition of Desiccation Tolerance and Longevity in Seeds of Arabidopsis thaliana (A Comparative Study Using Abscisic Acid-Insensitive abi3 Mutants). Plant Physiology 102, 1185–1191.

Oracz K, Bouteau HE-M, Farrant JM, Cooper K, Belghazi M, Job C, Job D, Corbineau F, Bailly C. 2007. ROS production and protein oxidation as a novel mechanism for seed dormancy alleviation. The Plant Journal 50, 452–465.

Pettigrew W, Dowd M. 2011. Varying Planting Dates or Irrigation Regimes Alters Cottonseed Composition. Crop Science 51, 2155.

Qanmber G, Ali F, Lu L, Mo H, Ma S, Wang Z, Yang Z. 2019a. Identification of Histone H3 (HH3) Genes in Gossypium hirsutum Revealed Diverse Expression During Ovule Development and Stress Responses. Genes 10, 355.

Qanmber G, Liu J, Yu D, Liu Z, Lu L, Mo H, Ma S, Wang Z, Yang Z. 2019b. Genome-Wide Identification and Characterization of the PERK Gene Family in Gossypium hirsutum Reveals Gene Duplication and Functional Divergence. International journal of molecular sciences 20, 1750.

Qanmber G, Lu L, Liu Z, Yu D, Zhou K, Huo P, Li F, Yang Z. 2019c. Genome-wide identification of GhAAI genes reveals that GhAAI66 triggers a phase transition to induce early flowering. Journal of Experimental Botany.

Qanmber G, Yu D, Li J, Wang L, Ma S, Lu L, Yang Z, Li F. 2018. Genome-wide identification and expression analysis of Gossypium RING-H2 finger E3 ligase genes revealed their roles in fiber development, and phytohormone and abiotic stress responses. Journal of Cotton Research 1, 1.

Rajjou L, Lovigny Y, Groot SPC, Belghazi M, Job C, Job D. 2008. Proteome-Wide Characterization of Seed Aging in Arabidopsis: A Comparison between Artificial and Natural Aging Protocols. Plant Physiology 148, 620–641.

Ravindran P, Verma V, Stamm P, Kumar PP. 2017. A Novel RGL2-DOF6 Complex Contributes to Primary Seed Dormancy in Arabidopsis thaliana by Regulating a GATA Transcription Factor. Molecular Plant 10, 1307–1320.

Razem FA, Baron K, Hill RD. 2006. Turning on gibberellin and abscisic acid signaling. Current Opinion in Plant Biology 9, 454–459.

Sakuma Y, Liu Q, Dubouzet JG, Abe H, Shinozaki K, Yamaguchi-Shinozaki K. 2002. DNA-Binding Specificity of the ERF/AP2 Domain of Arabidopsis DREBs, Transcription Factors Involved in Dehydration- and Cold-Inducible Gene Expression. Biochemical and Biophysical Research Communications 290, 998–1009.

Sano N, Rajjou L, North HM, Debeaujon I, Marion-Poll A, Seo M. 2016. Staying Alive: Molecular Aspects of Seed Longevity. Plant & amp; cell physiology 57, 660–674.

Shapiguzov A, Vainonen JP, Hunter K, Tossavainen H, Tiwari A, Järvi S, Hellman M, Aarabi F, Alseekh S, Wybouw B, Van Der Kelen K, Nikkanen L, Krasensky-Wrzaczek J, Sipari N, Keinänen M, Tyystjärvi E, Rintamäki E, De Rybel B, Salojärvi J, Van Breusegem F, Fernie AR, Brosché M, Permi P, Aro E-M, Wrzaczek M, Kangasjärvi J. 2019. Arabidopsis RCD1 coordinates chloroplast and mitochondrial functions through interaction with ANAC transcription factors. eLife 8, e43284.

Shu K, Zhang H, Wang S, Chen M, Wu Y, Tang S, Liu C, Feng Y, Cao X, Xie Q. 2013. ABI4 regulates primary seed dormancy by regulating the biogenesis of abscisic acid and gibberellins in arabidopsis. PLoS Genet 9, e1003577.

Smith CW. 1995. Crop production: evolution, history, and technology. New York: John Wiley and Sons.

Sugliani M, Rajjou L, Clerkx EJM, Koornneef M, Soppe WJJ. 2009. Natural modifiers of seed longevity in the Arabidopsis mutants abscisic acid insensitive3-5 (abi3-5) and leafy cotyledon1-3 (lec1-3). New Phytologist 184, 898–908.

Sun TP. 2008. Gibberellin metabolism, perception and signaling pathways in Arabidopsis. Arabidopsis Book 6, e0103.

Tejedor-Cano J, Prieto-Dapena P, Almoguera C, Carranco R, Hiratsu K, Ohme-Takagi M, Jordano J. 2010. Loss of function of the HSFA9 seed longevity program. Plant, Cell & Environment 33, 1408–1417.

Turley R, Chapman K. 2009. Ontogeny of Cotton Seeds: Gametogenesis, Embryogenesis, Germination, and Seedling Growth. 332–341.

Verma P, Kaur H, Petla BP, Rao V, Saxena SC, Majee M. 2013. PROTEIN L-ISOASPARTYL METHYLTRANSFERASE2 is differentially expressed in chickpea and enhances seed vigor and longevity by reducing abnormal isoaspartyl accumulation predominantly in seed nuclear proteins. Plant Physiology 161, 1141–1157.

Wan Q, Guan X, Yang N, Wu H, Pan M, Liu B, Fang L, Yang S, Hu Y, Ye W, Zhang H, Ma P, Chen J, Wang Q, Mei G, Cai C, Yang D, Wang J, Guo W, Zhang W, Chen X, Zhang T. 2016. Small interfering RNAs from bidirectional transcripts of GhMML3_A12 regulate cotton fiber development. New Phytologist 210, 1298–1310.

Wang Z, Cao H, Sun Y, Li X, Chen F, Carles A, Li Y, Ding M, Zhang C, Deng X, Soppe WJ, Liu YX. 2013. Arabidopsis paired amphipathic helix proteins SNL1 and SNL2 redundantly regulate primary seed dormancy via abscisic acid-ethylene antagonism mediated by histone deacetylation. The Plant Cell 25, 149–166.

Weigel D. 1995. The APETALA2 domain is related to a novel type of DNA binding domain. The Plant cell 7, 388–389.

Wu Z, Liang J, Zhang S, Zhang B, Zhao Q, Li G, Yang X, Wang C, He J, Yi M. 2018. A Canonical DREB2-Type Transcription Factor in Lily Is Post-translationally Regulated and Mediates Heat Stress Response. Frontier in Plant Science 9, 243.

Yamaguchi S, Kamiya Y, Nambara E. 2007. Regulation of ABA and GA Levels During Seed Development and Germination in Arabidopsis. 224–247.

Yang Z, Ge X, Yang Z, Qin W, Sun G. 2019. Extensive intraspecific gene order and gene structural variations in upland cotton cultivars. 10, 2989.

Yang Z, Zhang C, Yang X, Liu K, Wu Z, Zhang X, Zheng W, Xun Q, Liu C, Lu L, Qian Y, Xu Z, Li C, Li J, Li F. 2014. PAG1, a cotton brassinosteroid catabolism gene, modulates fiber elongation. New Phytologist 203, 437–448.

Zheng L, Wu H, Qanmber G, Ali F, Wang L, Liu Z, Yu D, Wang Q, Xu A, Yang Z. 2020. Genome-Wide Study of the GATL Gene Family in Gossypium hirsutum L. Reveals that GhGATL Genes Act on Pectin Synthesis to Regulate Plant Growth and Fiber Elongation. Genes (Basel) 11.

